# Gametophytic epigenetic regulators MEDEA and DEMETER synergistically suppress ectopic shoot formation in Arabidopsis

**DOI:** 10.1101/2023.05.16.541021

**Authors:** Mohit Rajabhoj, Sudev Sankar, Ramesh Bondada, Anju P Shanmukhan, Kalika Prasad, Ravi Maruthachalam

**Affiliations:** School of Biology, IISER-Thiruvananthapuram, Thiruvananthapuram, Kerala 695551, India; Department of Biology, IISER-Pune, Pune, Maharashtra 411008, India

**Keywords:** MEDEA, DEMETER, shoot development, epigenetic regulators, histone methylation, DNA methylation

## Abstract

The gametophyte to sporophyte transition facilitates the alternation of generations in a plant life cycle. The epigenetic regulators DEMETER(DME) and MEDEA(MEA) synergistically control central cell proliferation and differentiation, ensuring proper gametophyte to sporo-phyte transition in *Arabidopsis*. Mutant alleles of *DME* and *MEA* are female gametophyte lethal, eluding the recovery of recessive homozygotes to examine their role in the sporophyte. Here, we exploited the paternal transmission of these mutant alleles coupled with CENH3-haploid inducer to generate *mea*-1;*dme*-2 sporophytes. Strikingly, the simultaneous loss of function of MEA and DME leads to the emergence of ectopic shoot meristems at the apical pole of the plant body axis. *DME* and *MEA* are expressed in the developing shoot apex and regulate the expression of various shoot-promoting factors. Chromatin immunoprecipitation(ChIP), DNA methylation and gene expression analysis revealed several shoot regulators as potential targets of MEA and DME. Knockdown of upregulated shoot-promoting factors STM, CUC2, and PLT5 rescued the ectopic shoot phenotypes. Our findings reveal a previously unrecognized synergistic role of MEA and DME in restricting the meristematic activity at the shoot apex during sporophytic development to orchestrate the plant architecture.

## Introduction

Sporophyte development in higher plants results from a fine balance between cell proliferation and differentiation at the meristems. Primary meristems at the shoot and root apices are responsible for all post-embryonic organ development. This is coordinated by well guided distribution of plant hormones (Stahl and Simon 2010; Ten Hove et al. 2015). Throughout this process, epigenetic regulators provide an additional fine-tuning of gene expression, further contributing to plant developmental plasticity. Regulators of Histone methylation like Polycomb Repressive Complex 1 & 2 (PRC1 & PRC2) transcriptionally repress the expression of embryonic genes such as *BABYBOOM* (*BBM*), *LEAFY COTYLEDON1* & *2* (*LEC1* & *LEC2*) to control embryo development (Chen et al. 2010). Other epigenetic regulators such as EMBRYONIC FLOWER (EMF)- and VERNALIZATION (VRN)-PRC2s regulate sporophyte development during the transition from the vegetative to the reproductive life phase (Yoshida et al. 2001; De Lucia et al. 2008). Genes expressed during shoot development like *WUSCHEL* (*WUS*), *SHOOT MERISTEMLESS* (*STM*), *AGAMOUS* (*AG*), *BREVIPEDICELLUS* (*BP*), *KNOTTED*-*LIKE FROM ARABIDOPSIS THALIANA 2* (*KNAT2*), and *KNUCKLES* (*KNU*) are known targets of PRC1 and PRC2 components (Xu and Shen 2008; Chen et al. 2016). Their expression pattern is regulated by trimethylation of histone H3 at Lys27 (H3K27me3) repression mark (Kwon et al. 2006; Lodha et al. 2013; Sun et al. 2014; Xiao and Wagner 2015; Xiao et al. 2017). Other plants specific transcription factors such as *PLETHORA* (*PLT*) are also epigenetically regulated (Aichinger et al. 2011; Lee and Seo 2018). *PLT* genes control both root and shoot development (Prasad et al. 2011; Hofhuis et al. 2013). *PLT* genes activate the shoot-promoting factor *CUP-SHAPED COTYLEDON2* (*CUC2*) during de novo shoot meristem formation (Kareem et al. 2015). CUCs and STM positively regulate each other to maintain stem cell populations and promote organ formation at the shoot apical meristem (SAM) boundary (Spinelli et al. 2011). CUCs are also positively regulated by chromatin remodeling factors like BRAHMA (BRM), which is required to establish leaf primordia (Shen and Xu 2009). These studies indicate the maintenance of a delicate balance of CUCs expression during shoot development *via* direct or indirect epigenetic regulators. Apart from controlling shoot development, epigenetic marks such as H3K27me3 and DNA methylation are often reset during major developmental phase transitions, such as gamete and embryo formation (Slotkin et al. 2009; Iwasaki and Paszkowski 2014; Borg et al. 2020). Such resets facilitate transcription of array of genes in companion cells in gametophyte and floral transition genes in meristem essential to carry out the developmental phase change (Calarco et al. 2012; Jullien et al. 2012; You et al. 2017; Borg et al. 2020). Apart from genes, transposable elements (TEs) are also essential for successful developmental phase change (Slotkin et al. 2009; Ibarra et al. 2012; Higo et al. 2020). In absence of DNA methylation and histone modification, TEs express in ectopic tissue and disrupt such phase change events (Lafos et al. 2011; Baubec et al. 2014).

In Arabidopsis, embryo development is regulated by the components of FERTILIZATION INDEPENDENT SEED-Polycomb Repressive Complex 2 (FIS-PRC2) and DNA glycosylase DEMETER (DME) (Xiao et al. 2003; Gehring et al. 2006). DME and members of FIS-PRC2 complex like MEDEA (MEA) are primarily expressed in central cells of the embryo sac (Luo et al. 2000; Choi et al. 2002). The loss-of-function mutations in these genes result in female gametophyte lethality reflected as parent-of-origin defects during seed development. DME and MEA are known to regulate seed development together, where DME activates the expression of *MEA* in the central cell by removing methylated cytosines (mC’s) from *MEA* promoter and intergenic regions (Xiao et al. 2003; Gehring et al. 2006). DME is expressed in vegetative cell of male gametophyte and it is essential for male gametophyte function (Schoft *et al*. 2011; Park *et al*. 2017, Park *et al*. 2020).

Apart from gametogenesis and seed development, histone methylation mediated by PRC2 also affects seedling transitions by modulating seed dormancy and repression of embryonic traits in developing seedlings (Bouyer et al. 2011). Similarly, DNA methylation is reported to be high in developing seedlings to suppress transposon element expression and maintain an undifferentiated stem cell niche (Bouyer et al. 2017; Higo et al. 2020). These studies indicate an essential role of DNA and histone methylation in regulating the plant architecture. PRC2 members like MEA and DNA glycosylases like DME are likely to act in transcriptional regulation of target genes. Thus, a complete loss-of-function of *DME* and *MEA* provides an ideal system for examining their function during sporophytic development.

Mutant alleles of *DME* and *MEA* are transmitted through male gamete as microsporogenesis and microgametogenesis remain unaffected in most of these mutants, although *dme* and *mea* are not transmitted through female lineage (Grossniklaus et al. 1998; Choi et al. 2002; Köhler et al. 2003). The generation of homozygous gametophytic lethal mutants thus is challenged by conventional genetic approaches, which rely on self-pollination (inbreeding) of heterozygous single/double mutants to generate respective homozygotes barring those alleles with the leaky transmission. Hence, it is difficult to decipher the role of such gametophyte-lethal genes, if any, in sporophytic development. Studies involving maternal lethals of *DME*, *MEA,* and their epigenetic regulation in sporophytes thus relied either on examining heterozygotes (Roszak and Köhler 2011; Schoft et al. 2011; Figueiredo et al. 2015; Park et al. 2017; Zhang et al. 2018; Khouider et al. 2021) or homozygous plants carrying a weak allele (Choi et al. 2002; Park et al. 2017; Simonini et al. 2021; Kim et al. 2021; Williams et al. 2022), thereby restricting complete understanding of their function. Alternative methods to study DME and MEA in sporophyte development have been employed previously. These include paternal haploids from single mutants, conditional homozygotes, and homozygotic progeny of transheterozygote mutants (Ravi et al. 2014; Roy et al. 2018; Simonini et al. 2021; Kim et al. 2021).

Here, we employed haploid genetics to generate viable *mea*^-/-^;*dme*^-/-^ doubled haploid (DH) sporophytes. Using these DH mutant homozygotes, we show that MEA and DME are essential for sporophytic development in *Arabidopsis*. Loss of function of *MEA* and *DME* leads to initiation of ectopic shoot meristems at the apical pole of the plant body axis. We further show that these epigenetic regulators control shoot meristem development by repressing key shoot-promoting factors such as *STM*, *PLT5*, and *CUC2*. Our studies reveal a new role of MEA and DME mediated epigenetic regulation in controlling shoot architecture during postembryonic development.

## Results

### *mea*-1^-/-^;*dme*-2^-/-^ double mutants are viable and initiate functional ectopic SAM

Previous studies have established that MEA and DME play a key role in seed development by regulating polar nuclei and endosperm division after double fertilization (Grossniklaus et al. 1998; Choi et al. 2002). Several *dme* and *mea* alleles have been isolated by induced mutagenesis from a few select natural *Arabidopsis thaliana* accessions (Choi et al. 2002; Guitton et al. 2004; Schoft et al. 2011). Irrespective of the accession, both *dme* and *mea* mutant alleles can be transmitted through pollen (male) with rare or no maternal female transmission resulting in >25% seed lethality upon self-fertilization of the corresponding heterozygotes. We have earlier shown that generating *mea* and *dme* paternal haploid plants is possible using CENH3-mediated haploid induction (Ravi *et al*. 2014; Roy *et al*. 2018; Supplementary Fig. S1). Here, we advanced further to generate doubled haploid (DH) plants of *dme*-2^-/-^ (L*er*), *dme*-6^-/-^, *dme*-7^-/-^(Col-0), and *mea*-1^-/-^ (L*er*), *mea*-6^-/-^, *mea*-7^-/-^ (C24) mutants to examine the recessive vegetative phenotypes associated with *dme* and *mea* mutants (Figure 1A; Supplementary Table 1). Here, we used *dme*-2 and *mea*-1 alleles for generating double homozygotes for the following reasons; firstly, among all the mutant alleles characterized, *mea*-1 and *dme*-2 are reported to show strong loss-of-function phenotypes (Grossniklaus et al. 1998; Choi et al. 2002; Simonini et al. 2021; Kim et al. 2021), which is ideal for double mutant studies. Secondly, both alleles are of the Landsberg *erecta* (L*er*) ecotype, which yields a higher number of spontaneous DH seeds than Columbia (Col-0) (Ravi and Chan 2010). This feature is essential to maximize seed yield to obtain viable spontaneous doubled haploid seeds in the strong gametophyte lethal alleles with high seed abortion like *mea-*1 and *dme-*2. Thirdly, identical L*er* backgrounds of single mutants of *mea* and *dme* ensure genetic homogeneity in the resultant double mutants, thereby avoiding confounding effects due to different genetic backgrounds. We could not generate double mutants of *dme* and *mea* alleles in Col-0 and C24 due to severe seed lethality in these backgrounds (Supplementary Table 1).

**Fig. 1.**
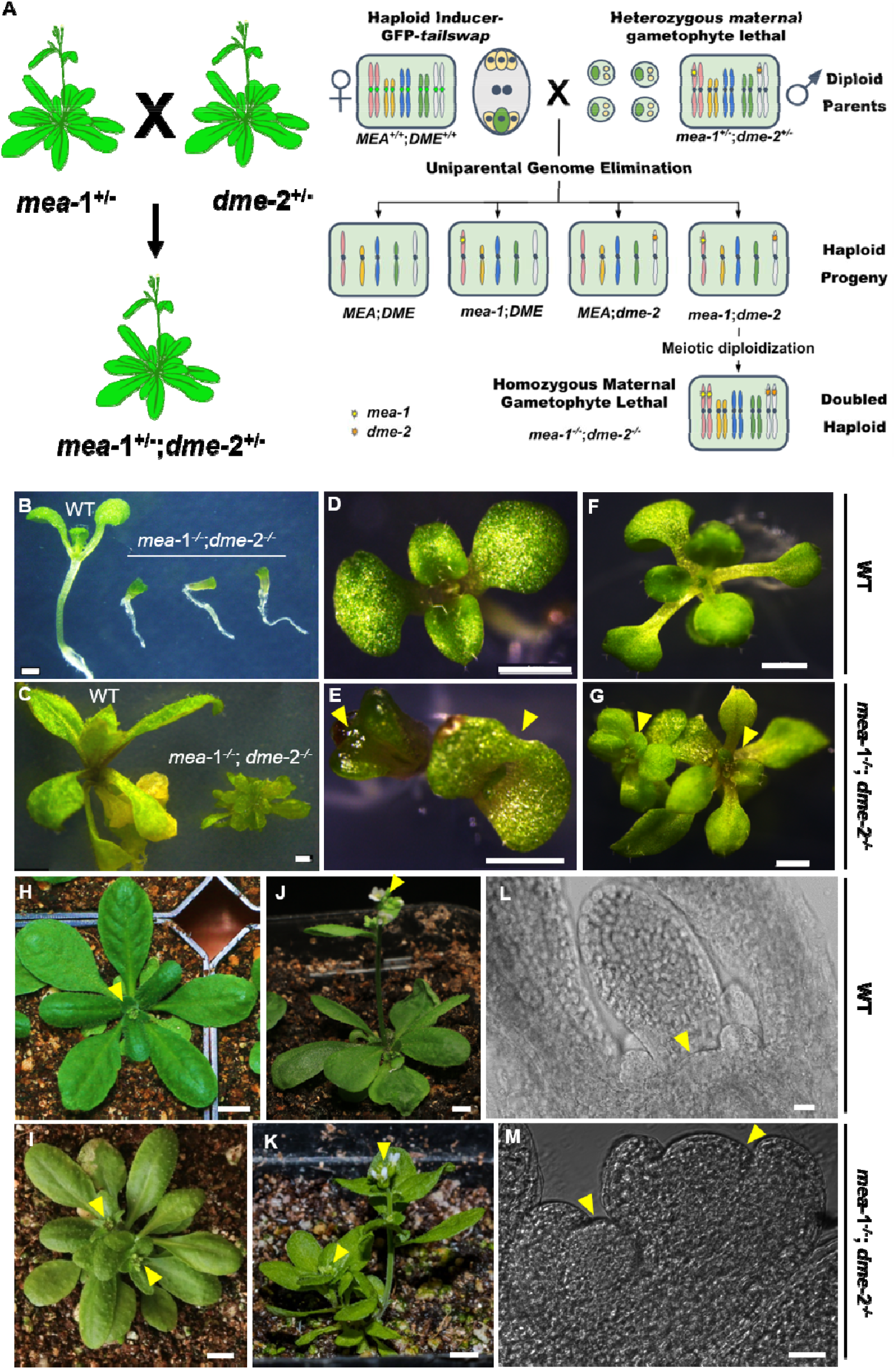
Generation and phenotypic examination of *mea*-1-/-;*dme*-2-/-plants. **(A)** Schematic diagram illustrating rescue of *mea*-1-/-;*dme*-2-/-plants using CENH3-mediated genome elimination. **(B,C)** *mea*-1-/-;*dme*-2-/-seedlings as compared to WT at 2-leaf (5dpg) and 6-leaf (10 dpg) stage respectively. Representative images of L*er* WT plants at **(D)** 2-leaf (5dpg), **(F)** 6-leaf (10 dpg), **(H)** inflorescence emergence (21 dpg) and **(J)** flowering (28 dpg) stage as compared with *mea*-1-/-;*dme*-2-/-plants with ectopic shoot (marked by yellow arrowheads) at **(E)** 2-leaf, **(G)** 6-leaf, **(I)** inflorescence emergence and **(K)** flowering stages. Longitudinal section of shoot apex of the 3-leaf stage (7 dpg) **(L)** L*er* WT and **(M)** *mea*-1-/-;*dme*-2-/-seed-ling. Yellow arrowhead is pointing at the shoot apex. Scale bar represents **(B-G)**= 1 mm; **(H-K)**= 5 mm; **(L-M)**= 20 µm. dpg-days post germination.

**Table 1.** Context-based cytosine methylation associated with genes and transposable elements (TEs). Percent of context-based methylation is calculated using Bismark methylation extraction algorithm.

| Context | WT | *mea-1<sup>-/-</sup>;dme-2<sup>-/-</sup>* |  |
| --- | --- | --- | --- |
|  |  | Associated with genes | Associated with TEs |
| mC's in CpG context, % | 15463967, 22.50% | 26006159, 24.60% | 392421, 6.64% |
| mC's in CHG context, % | 6676430, 9.80% | 11864625, 11.10% | 465643, 7.88% |
| mC's in CHH context, % | 13466882, 4.40% | 22075839, 4.50% | 1729891, 29.28% |
| Total mC's, % | 35607279, 36.70% | 59946623, 40.20% | 2587955, 43.81% |

*mea*-1 haploid seedlings (n=14) did not show any visible phenotypic defects and grew normal like wild-type (WT). *mea*-1 haploids yielded spontaneous DH seeds, of which 2% (n= 21/1178; Supplementary Table 1) were viable, giving rise to *mea*-1^-/-^ DH offspring. Consistent with previous studies, most seeds produced by DH *mea*-1^-/-^ plants were atrophied (81%, n= 1407/1754; Supplementary Fig. S1) (Simonini et al. 2021). A small fraction of germinated seedlings (4%, n= 14/338) failed to grow two to four days post-germination (dpg). Such seedlings only developed cotyledons and a pair of young leaves. Unlike *mea*-1 haploids, a fraction of *mea*-1^-/-^ (33%, n= 107/324) displayed stunted and aberrant vegetative growth (Supplementary Fig. S2). However, when transferred to the soil, these seedlings recovered and showed normal vegetative growth with no discernible phenotypes.

Like *mea*-1 haploids, *dme*-2 haploids (n= 15) also showed normal vegetative growth but severe seed abortion during reproductive development (98%, n= 1286/1308), owing to the strong loss-of-function of *dme*-2. Spontaneous *dme*-2^-/-^ DH seeds harvested from *dme*-2 haploid plants showed less than 2% germination (n= 22/1308; Supplementary Table 1). About 10% (n= 43/478) of *dme*-2^-/-^ seedlings ceased to grow two to four days post-germination (dpg) (Supplementary Fig. S2). Unlike *mea*-1^-/-^ seedlings, a vast majority of *dme*-2^-/-^ seed-lings (78%; n= 341/435) exhibited developmental abnormalities such as fused cotyledons, delayed emergence of true leaves, stunted hypocotyl, and retarded root growth (Supplementary Fig. S2). Despite these abnormalities, most of the *dme*-2^-/-^ seedlings (n= 153/157) recovered from the early vegetative growth defects when they transitioned to the adult stage, similar to that observed for *mea*-1^-/-^ homozygotes. However, unlike *mea*-1^-/-^ plants, *dme*-2^-/-^ plants displayed floral defects like altered petal number and positioning, and abnormalities in gynoecium, as reported in earlier studies (Choi *et al*. 2002; Ravi *et al*. 2014; Kim *et al*. 2021).

Next, we generated *mea*-1^-/-^;*dme*-2^-/-^ double homozygotes to examine the loss of histone methylation (H3K27me3) and DNA demethylation mediated by MEA and DME, respectively, during early vegetative development of seedlings. To this end, we crossed *mea*-1^+/-^;*dme*-2^+/-^ as a pollen parent, with haploid inducer GFP-*tailswap* as female, and selected *mea*-1;*dme*-2 haploids (n=9/32) and *MEA*;*DME* haploids (n=10/32) by PCR-based genotyping (Figure 1A; Supplementary Fig. S1). We collected spontaneous DH seeds from both the haploid sibling genotypes to generate diploid(=DH) *mea*-1^-/-^;*dme*-2^-/-^ and *MEA^+^*^/*+*^;*DME^+^*^/*+*^ plants. Such *MEA^+^*^/*+*^;*DME^+^*^/*+*^ DH plants were used as WT control in all our experiments involving *mea*-1^-/-^;*dme*-2^-/-^ mutants.

The DH seeds obtained from *mea*-1;*dme*-2 haploid plants showed less than 1% (n= 13/1950; Supplementary Table 1) viability up to germination (Supplementary Fig. S1). The *mea*-1;*dme*-2 haploids showed normal vegetative development like *mea*-1 and *dme*-2 haploids. The selfed *mea*-1^-/-^;*dme*-2^-/-^ plants displayed floral defects (Supplementary Fig. S3) and continued to produce atrophied seeds (n= 1678/1681), as observed in the parent generation. Around 38% (n= 87/226) of germinated *mea*-1^-/-^;*dme*-2^-/-^ were arrested in development and displayed pale and fused cotyledons (Supplementary Fig. S2). The remaining viable *mea*-1^-/-^;*dme*-2^-/-^ seedlings displayed severely stunted growth and shorter hypocotyl compared to WT (Figure 1B). Such seedlings also showed underdeveloped and fused cotyledons without petioles compared to WT (Figure 1C-E). We also observed smaller, asymmetric true leaves, indicating defects during shoot development (n= 107/139). Remarkably, nearly half the number of *mea*-1^-/-^;*dme*-2^-/-^ seedlings (n= 71/139) displayed ectopic shoot emerging from the existing shoot apex (Figure 1F, G). Further analysis revealed the initiation of ectopic shoot meristems in *mea*-1^-/-^;*dme*-2^-/-^ (n= 25/48). Figure 1L, M). It may be argued that the ectopic shoot phenotype is a manifestation of aneuploidy resulting from haploid induction crosses (Tan et al. 2015). However, we rule out this possibility because rarely such aneuploidy syndromes show such prominent ectopic shoot phenotypes. In addition, we carried out chromosome dosage analysis from whole genome bisulfite sequencing reads to confirm that such plants with ectopic shoot phenotype are diploids (Supplementary Fig. S1). Furthermore, we consistently observed double shoot phenotype from plants derived from 9 independent *mea*-1;*dme*-2 haploids(biological replicates) further ruling out aneuploidy as the cause of the double shoot phenotype.

Like *mea*-1^-/-^ and *dme*-2^-/-^ single mutant plants, *mea*-1^-/-^;*dme*-2^-/-^ plants also recovered from their early developmental defects and displayed normal vegetative growth in the later stages (n= 129/132, Figure 1I; Supplementary Fig. S3). In these *mea*-1^-/-^;*dme*-2^-/-^ plants two shoot meristems developed independently and upon transition to reproductive phase, they gave rise to two inflorescence shoots from a single vegetative body (rosette) (Figure 1I, K; Supplementary Fig. S3). This “twin-plant” phenotype is reminiscent of conjoined twins having two independent upper trunks with a common lower portion of the body. The “twin plant” phenotype arising from persistent ectopic shoots suggests a role for DME and MEA in suppressing the initiation of ectopic shoot meristems at apical pole of plant body axis.

Interestingly, a fraction of single mutant *mea*-1 (n= 217/324), *dme*-2 (n= 94/435), and *mea*-1;*dme*-2 double mutants (n=32/139) did not show any visible developmental defects and grew like WT. Even single and double mutant seedlings that show early developmental defects post-germination show resilience at later stages of vegetative development. However, the ectopic shoots initiated during early developmental stages persist throughout the plant life cycle, displaying twin plant phenotype. This observation suggests that the function of both DME and MEA is predominantly required only during juvenile growth but dispensable for later vegetative growth.

### MEA and DME are expressed in developing shoot meristems and leaf primordia

Earlier studies have shown that in addition to its gametophyte expression in the central cell of the female gametophyte and in the vegetative cell of the male gametophyte, *DME* is also expressed in sporophytic tissues such as the developing SAMs, root apical meristems (RAMs) and inflorescence meristems (IMs) (Kim *et al*. 2008, Kim *et al*. 2021). Likewise, *MEA* is expressed in the developing embryo till the torpedo stage (Vielle-Calzada et al. 1999; Spillane et al. 2007) and induced in sporophytic tissues on demand when challenged with pathogens (Roy et al. 2018). Further, transcriptome data also indicates the expression of MEA and DME mRNA in the seedlings and other sporophytic tissues (Schmid et al. 2005). However, spatio-temporal expression patterns of *MEA* and *DME* in the shoot meristems remained largely unknown. Since we observed ectopic SAM initiation in *mea-*1^-/-^;*dme-*2^-/-^ seedlings, we examined if MEA and DME are expressed in shoot meristem. We first monitored the transcript levels of *MEA* and *DME* in growing seedlings by RT-qPCR*. DME* transcript levels increased in 2 dpg seedlings, with a decline in expression after 3 dpg. Its transcript levels increased after 5 dpg with plant growth (Figure 2A). On the other hand, we observed *MEA* mRNA expression to be low in the early stage of vegetative development but steadily increased over time till the 10 dpg seedling stage (Figure 2B).

**Fig. 2.**
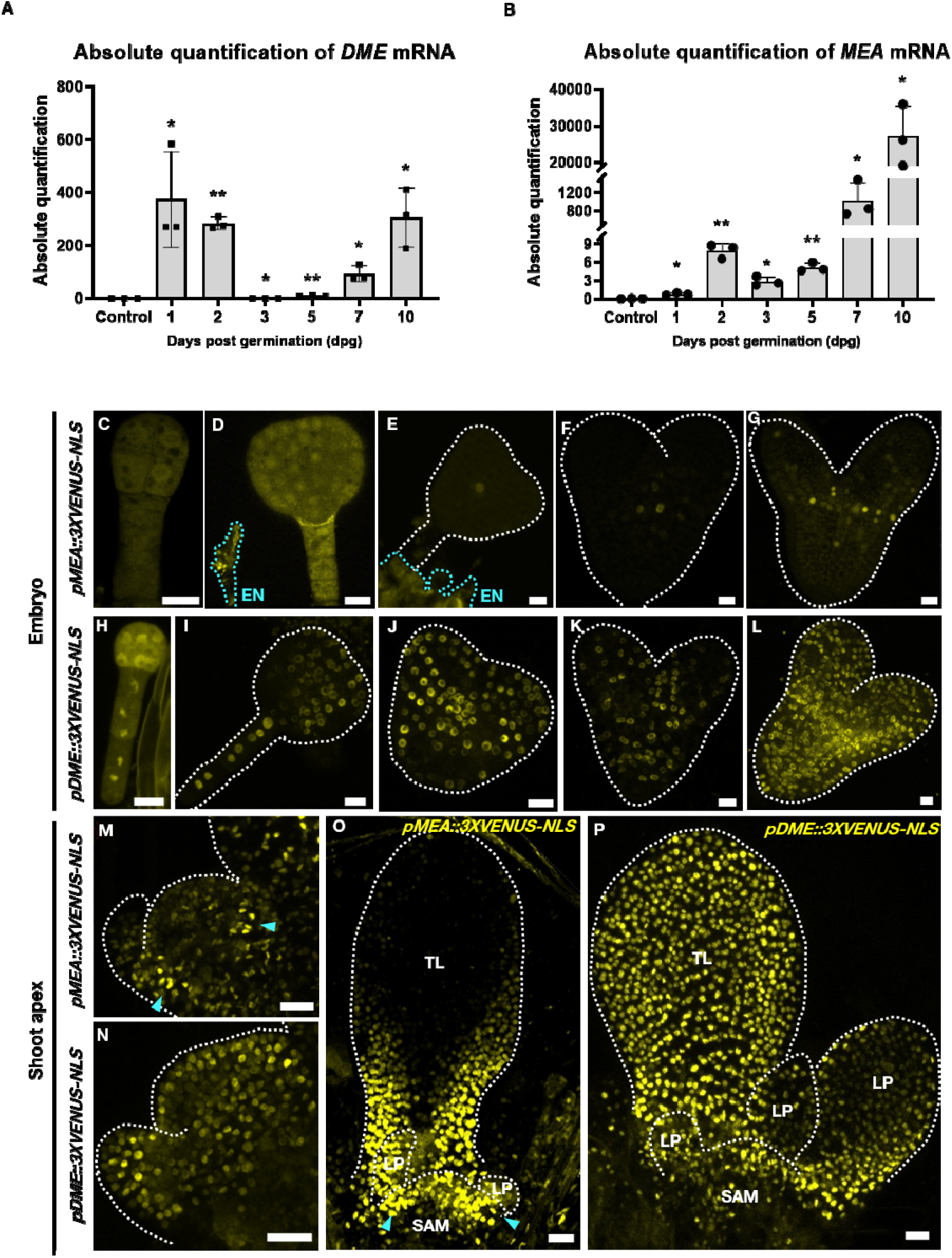
Expression analysis of MEA and DME in sporophytic tissues (A,B) Relative abundance of DME and MEA mRNA in developing seedlings. **(A)** *DME* mRNA levels are abundant in 1-and 2-dpg seedlings with sharp decrease in its level on 3-and 5-dpg (1-dpg: 373.62±181.9, \**p*= 0.0407; 2-dpg: 282.80±24.20, \*\**p*= 0.0024; 3-dpg: 0.61±0.13, \**p*= 0.138; 5-dpg: 9.92±1.63, \*\**p*= 0.0089; 7-dpg: 92.54±30.44, \**p*=0.0342; 10-dpg: 304.79±112.62, \**p*= 0.0426). **(B)** *MEA* mRNA levels gradually increased as the seedling transitioned from 1- to 10-dpg (1-dpg: 0.76±0.21, \**p*= 0.035; 2-dpg: 7.88±1.19, \*\**p*= 0.0076; 3-dpg: 2.83±0.77, \**p*= 0.024; 5-dpg: 5.14±0.76, \*\**p*= 0.0076; 7-dpg: 1020.56±404.7, \**p*=0.0486; 10-dpg: 27055.34±8463.32, \**p*= 0.031). Statistical test: Multiple unpaired *t*-tests (Holm-Sidak method), with alpha = 0.05. Error bar represents SD. **(C-P)** Representative images of embryos and shoot apices showing expression patterns of **(C-G, M,O)** *pMEA*::*3XVENUS-NLS* and **(H-L, N,P)** *pDME*::*3XVENUS-NLS. pMEA*::*3XVENUS-NLS* expression (yellow) in developing embryo **(C)** octant, **(D)** globular, **(E)** early heart-shape, **(F,G)** heart-shape **(M)** SAM and in **(O)** shoot apex along with peripheral domain of SAM marked by blue arrowhead. *pDME*::*3XVENUS-NLS* expression (yellow) in developing embryo **(H)** octant, **(I)** globular **(J)** early heart-shape, **(K,L)** heart-shape **(N)** SAM and **(P)** shoot apex. In panels **D**, **E** residual endosperm tissues also show signal (outlined by blue dotted lines) along with embryo. EN-Endosperm, LP-leaf primordium, SAM-shoot apical meristem, TL-true leaf. Scale bar represents **(C-L)**= 10 µm; **(M-P)**= 20 µm.

Next, we examined the spatiotemporal expression pattern of DME and MEA using confocal based imaging of the WT plants harboring *pMEA*::*gMEA*:*vYFP* and *pDME*::*gDME*:*vYFP* translational fusions. Both, *pMEA*::*gMEA*:*vYFP* and *pDME*::*gDME*:*vYFP* transgenes were functional and rescued the *mea*-1^-/-^ and *dme*-2^-/-^ mutant phenotypes (Supplementary Fig. S4; Supplementary Table 1). The rescue in seed lethality and expression pattern suggests that both the transgenes harbor necessary promoter sequences capable of recapitulating the endogenous expression pattern of *MEA* and *DME*. Consistent with the previous studies, we observed active *MEA* transcription in developing embryos (Vielle-Calzada et al. 1999; Spillane et al. 2007). We observed *pMEA*::*3XVENUS-NLS*(nuclear localization signal) activity during globular stage of embryo. The expression was weak, dispersed and could be detected only in a few cells during later stages of embryo development (n= 33/45) (Figure 2C-G). Unlike *pMEA*::*3XVENUS-NLS*, nearly uniform expression of *pDME*::*3XVENUS-NLS* was observed throughout the embryo development, right from the four celled to the walking-stick stage (n= 66/71) (Figure 2H-L).

Interestingly, during post-embryonic development. *MEA* was predominantly expressed in the peripheral domain of the shoot meristem (Figure 2M). Its expression was also detected in developing leaf primordia (Figure 2O). As reported earlier, *DME* was expressed uniformly throughout the shoot meristem (Figure 2N) (Kim *et al*. 2008, Kim *et al*. 2021). Like *MEA*, its expression was also detected in developing leaf primordia (Figure 2P).

### MEA and DME repress the expression of shoot-promoting factors

We next examined if the ectopic shoot meristems in *mea-*1^-/-^;*dme-*2^-/-^ made functional stem cell niches. *WUSCHEL* is essential for maintenance of the shoot stem cell niche (Mayer et al. 1998; Schoof et al. 2000). We therefore monitored the expression of *pWUS*::*3XVENUS-NLS*:*tWUS*, a reporter which specifically marks the shoot stem cell organizer (Varapparambath et al. 2022). Strikingly, *mea-*1^-/-^;*dme-*2^-/-^ seedlings showed ectopic *WUS* promoter activity at the shoot apex (n= 21/48, Figure 3C-D, F, F(i) to F (iii)) as compared to WT seedlings where reporter activity was confined to one organizing center (Figure 3E). Ectopic WUS expression corroborates with shoot phenotype seen in *mea*-1^-/-^;*dme*-2^-/-^. We also observed ectopic WUS-YFP activity in developing cotyledons of *mea*-1^-/-^;*dme*-2^-/-^ embryos and SAM (n= 26/78) indicating aberrant mixed cell fate in the mutant embryo (Figure 3C-D,F). This indicates de novo organizing centers specified in the non-canonical regions in the embryos, as reported earlier (Simonini et al. 2021).

**Fig. 3.**
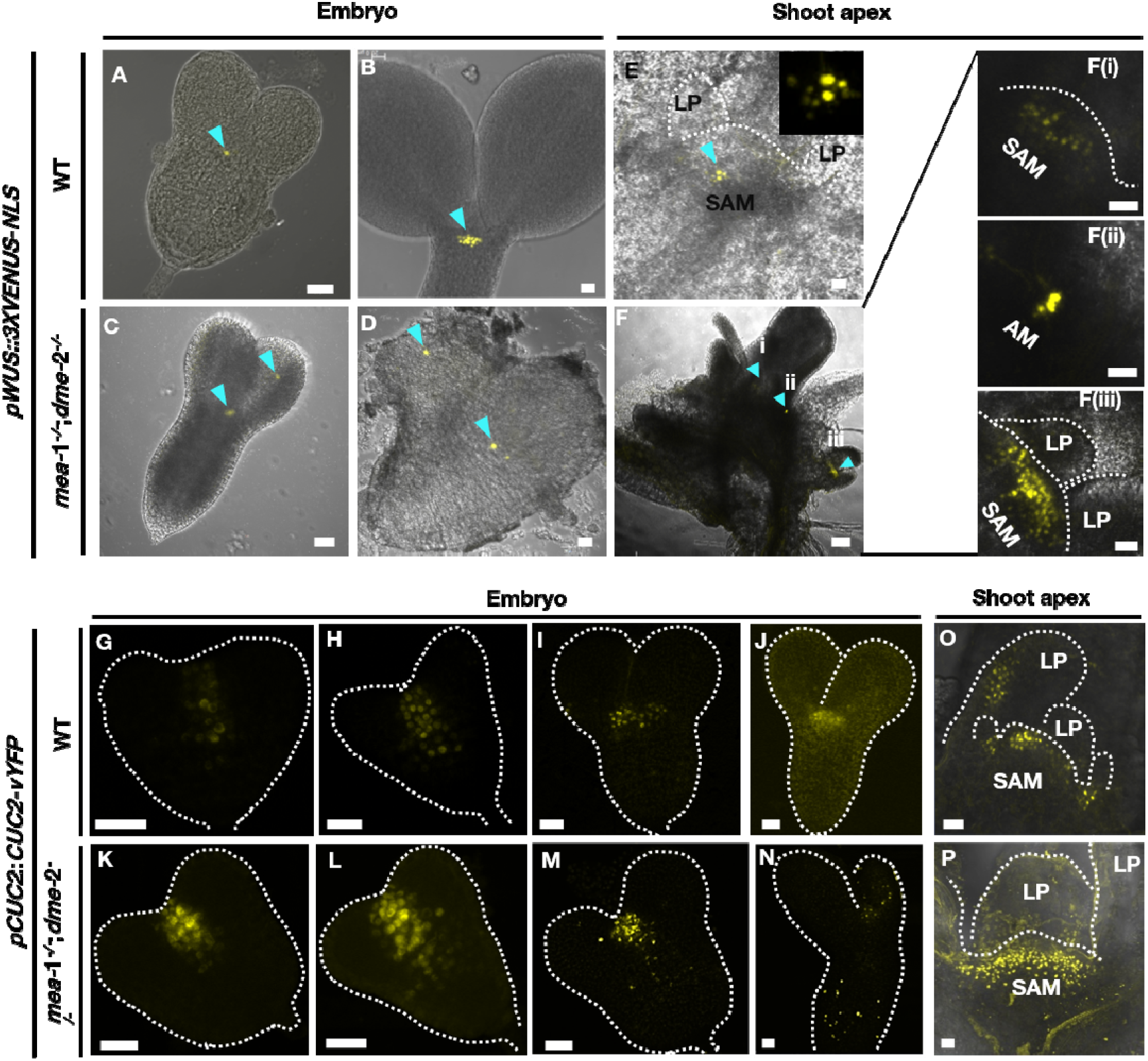
Deregulated expression pattern of shoot regulators in *mea-*1*-/-*;*dme-*2*-/-* embryos and seedling (A-D’) Representative images of embryos and shoot apices showing expression pattern of **(A-F(iii))** *pWUS*::*3XVENUS-NLS,* **(G-P)** *pCUC2*::*CUC2-vYFP* shoot regulators. *pWUS*::*3XVENUS-NLS* expression (yellow) in organizing center of embryos of **(A-B)** wild-type and **(C-D)** *mea-*1*-/-*;*dme-*2*-/-(*n= 26/78) showing ectopic organizing center in cotyledon (marked with yellow arrowhead). *pWUS*::*3XVENUS-NLS* is marking center of shoot meristem in **(E)** wild-type, inset is showing organizing center in SAM and **(F to F(iii))** *mea-*1*-/-*;*dme-*2*-/-,*along with **(F(i))** primary shoot meristem **(F(ii))** axillary meristem and **(F(iii))** ectopic shoot meristem, marked with blue arrowhead (n=21/48). The meristem dome (SAM) and leaf primordia (LP) are labeled and indicated with a white dashed line. Scale bar represents 20 µm in **(A-E)** and **(F(i) to F(iii))**, and 100 µm in **(F)**. **(G-P)** *pCUC2*::*CUC2-vYFP* (yellow) expression in embryos of **(G-J)** wild-type and **(K-N)** *mea-*1*-/-*;*dme-*2*-/-* showing ectopic expression at SAM and developing cotyledons (n= 31/77). *pCUC2*::*CUC2-vYFP* expression in shoot apex of **(O)** wild-type and **(P)** *mea-*1*-/-*;*dme-*2*-/-* showing ectopic expression in SAM apart from leaf primordia (n= 15/77). SAM-shoot apical meristem, AM-axillary meristem, LP-leaf primordium. Scale bar represents 20 µm.

We further analyzed the expression of genes that regulate the boundary between the meristem and developing lateral organ primordia. The transcription factor, CUC2, is an early marker of shoot organ boundary specification at SAM (Heisler et al. 2005). CUC2 expression in developing embryo displays characteristic stripe formation in the central domain away from protoderm cells till the heart shape stage but later surrounds the SAM between its boundary with cotyledons (Aida et al. 1999, 2002) (Figure 3G-J). We monitored CUC2 expression by following *pCUC2*::*CUC2*-*vYFP*; reporter in *mea*-1^-/-^;*dme*-2^-/-.^. The typical pattern of *pCUC2*::*CUC2*-*vYFP* in the central domain of embryo was lost. Rather, we observed dispersed expression in SAM initial and protoderm cells (n= 31/77). (Figure 3K,L). We observed *pCUC2*::*CUC2*-*vYFP* expression in the cotyledon primordia of a few embryos, further suggesting mixed cell fate identity in the mutant embryos (n=15/77) (Figure 3M,N). Interestingly, *pCUC2*::*CUC2*-*vYFP* expression in *mea-*1^-/-^;*dme-*2^-/-^ shoots is dispersed along the peripheral and central zone instead of being restricted at the organ primordia (Figure 3P). The ectopic expression of *pCUC2*::*CUC2*-*vYFP* indicates the de-repression of *CUC2* in SAM. These data demonstrate the de-repression of most canonical shoot regulators in *mea-*1^-/-^;*dme-* 2^-/-^ seedlings.

### Shoot regulators are targets of MEA and DME during early shoot development

The ectopic shoot phenotype in *mea*-1^-/-^;*dme*-2^-/-^ plants and the expression of *DME* and *MEA* in the shoot meristem, prompted us to identify critical targets of MEA and DME that regulate the shoot meristem specification. The interplay between DNA methylation and histone modifications in specifying cell fate and differentiation during development is well-known in animals and plants (Du et al. 2015; Williams and Gehring 2020). For instance, a positive feedback loop between DNA methylation and H3K9 methylation maintains the heterochromatin in a repressed state in plants (Du et al. 2015). H3K27 methylation, similarly, has been observed predominantly in the heterochromatin state associated with non-CG methylation. The preferential binding of the histone demethylase RELATIVE OF EARLY FLOWERING 6 (REF6) to hypo-methylated DNA leads to the loss of the H3K27me3 mark from the chromatin (Cui et al. 2016; Qiu et al. 2019). Hence, we hypothesized that a similar interaction between DNA methylation and histone methylation could be jointly mediated by the action of DME and MEA, leading to the regulation of the shoot regulators.

As a histone methyltransferase, MEA catalyzes the trimethylation of H3K27. Hence, to identify targets of MEA, we performed whole-genome profiling of H3K27me3 levels by Chromatin Immunoprecipitation (ChIP)-seq experiment. Strikingly, we observed 4993 significant peaks in WT as compared to 4798 peaks in *mea-*1^-/-^;*dme-*2^-/-^ seedlings (Supplementary Table 2). In differential binding analysis we identified 91 genes with significant differential H3K27me3 binding. Apart from these 91 genes, we checked additional 3885 H3K27me3 binding loci and selected 431 enriched gene loci for further analysis based on their role in shoot development as predicted by the Gene Ontology Resource (http://geneontology.org/) (Figure 4A; Supplementary Table 2). These 431 genes encompass known shoot regulators like *STM*, *KNAT1*, *CUC2*, *PLT5*, *ESR1* & *2*, *CKX3*, *AHP6*, *KNU*, and *YUC4*. Upon closer examination of the broader peaks of H3K27me3, we found H3K27me3 binding more persistent in the promoter region (90.4%) of these genes, as reported by earlier studies (Zhang et al. 2007; Xiao et al. 2016). We also observed differential H3K27me3 binding in the gene body region of *CUC2*, *PLT5*, *STM*, and *WUS* (Figure 4B-E). This loss of in *mea-*1^-/-^;*dme-*2^-/-^ for such genes predicts their active transcription. Indeed, transcript levels of various shoot regulators which lost the repressive H3K27me3 mark such as STM, CUC2, PLT5, and WUS were increased in the mutant seedlings as compared to WT (Figure 3A-F, G-P, Supplementary Fig. S5). Overall, we observed a de-repression of various shoot regulators, in *mea-*1^-/-^;*dme-*2^-/-^ seedlings.

**Fig. 4.**
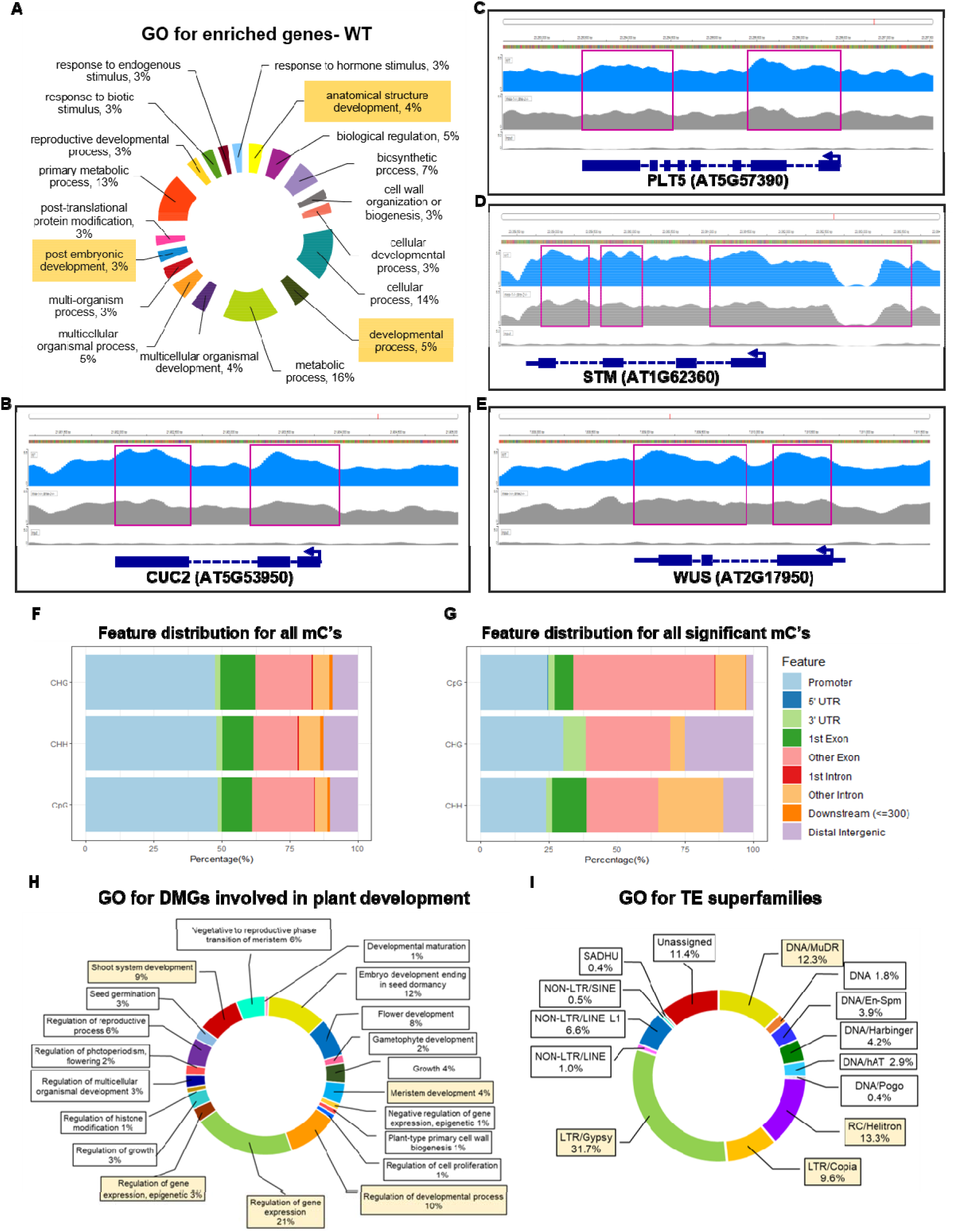
Validation of targets of MEA and DME (A) Gene ontology (GO) of enriched genes in ChIP using αH3K27me3. **(B-E)** Graphical representation of ChIP-seq using αH3K27me3 for **(B)** *STM*, **(C)** *CUC2* **(D)** *PLT5* and **(E)** *WUS* gene and their upstream region. Pink boxes denote prominent enrichment in WT ChIP DNA as compared to *mea-*1^-/-^;*dme-*2^-/-^ ChIP DNA. Input DNA is WT DNA without αH3K27me3 treatment. **(F-G)** Distribution of methylation locations at **(F)** across the genome and **(G)** at significant (≥25% methylation, *q*<0.01, *p*> 0.05) locations. **(H-I)** GO analysis for differentially methylated **(H)** genes (DMGs) involved in plant development **(I)** Transposable elements (TEs) classified in superfamilies.

To examine if loss of function of DME caused any change in methylation landscape of the *mea*-1^-/-^;*dme*-2^-/-^, we performed whole-genome bisulfite sequencing on *mea*-1^-/-^;*dme*-2^-/-^ and WT seedlings (Figure 4F-I, Supplementary Fig. S6). We observed a higher CpG and CHG methylation in *mea*-1^-/-^;*dme*-2^-/-^ seedlings (24.60% and 11.10%, Table 1) as compared to WT seedlings (22.50% and 9.80%, Table 1). However, the CHH methylation pattern remained unaffected. This suggests an increase in DNA methylation due to lack of functional DME in the *mea*-1^-/-^;*dme*-2^-/-^ plants. The total number of differentially methylated cytosines (DMCs) are significant (48.54% CpG) in the promoter region (2kb upstream of TSS) of annotated genes of *mea*-1^-/-^;*dme*-2^-/-^ seedlings. However, significant DMCs (more than 25% methylated) were found in the gene body region (72.26% CpG, 44.43% CHG, 65.21% CHH; Figure 4F,G), postulating a transcription activating role of methylation in *mea-*1^-/-^;*dme-*2^-/-^ plants. Gene Ontology analysis of genes associated with such DMCs show numerous genes involved in shoot system development like *A. thaliana RING finger 1A & 1B (AtRING1A* & *1B*), *Arabidopsis response regulator 1* & *12* (*ARR1* & *ARR12*), RNA-dependent RNA polymerase 6 (*RDR6*) (Figure 4H). For genes like *STM*, *CUC2*, *ESR1*, *ESR2*, *KNAT1* & *KNAT6*, we observed subtle hypermethylation in the promoter region (Supplementary Table 3). We also performed CHOP-PCR (Zhang et al. 2014; Dasgupta and Chaudhuri 2019) for detecting differentially methylated STM and WUS regulatory regions. We observed weak amplification for *WUS* and *STM* promoter regions in *mea*-1^-/-^;*dme*-2^-/-^ seedlings, indicating methylated DNA region (Supplementary Fig. S7). On the other hand, the gene body methylation pattern for such genes showed low levels of hypermethylation at both intronic and exonic cytosines in *mea-*1^-/-^;*dme-*2^-/-^. Although weak, these DNA methylation patterns for *STM*, *CUC2* and *WUS* are suggestive of aiding in the process of transcription, evident with their increase in expression levels in *mea-*1^-/-^;*dme-*2^-/-^ seedlings. DME is also known to positively regulate TEs in developing seeds and pollen grain. Apart from gametophyte, TEs are also differentially expressed in SAM, where they function in regulating genes like *AGO5*,*9* and shoot regulators like *STM*, *CUC2* and *CLV3* (Gutzat et al. 2020). We found a significant number of TEs (6278) to be differentially methylated in *mea*-1^-/-^;*dme*-2^-/-^ seedlings as compared to WT seedlings (Figure 4I). These TEs belong to Gypsy, MuDR, Helitron and COPIA superfamilies (Figure 4I, Supplementary Table 3) known to be specifically expressed in SAM during shoot development (Gutzat et al. 2020). Such change in methylation state of these TEs may also be associated with their activation or silencing, as seen during developmental phase change and after treatment with methylation inhibitor (Feng et al. 2010; Baubec et al. 2014). Here, we report an extension of function of DME in the early sporophytic tissue in regulating TEs expression and their subsequent function of silencing genes important in shoot development.

### Downregulation of shoot regulators in *mea-*1^-/-^;*dme-*2^-/-^ leads to a partial rescue of the ectopic shoot phenotype

Since the majority of shoot regulators are over-expressed in the SAM of *mea-*1^-/-^;*dme-*2 ^-/-^ seedlings (Supplementary Fig. S5), we sought if down-regulation of some of these shoot regulators viz., STM, PLT5, and CUC2 would suppress the ectopic shoot formation. Our selection of these genes for further analysis is based on their previously implicated role in promoting the shoot meristem formation. For instance, CUC2 shown to be sufficient for ectopic shoot meristem initiation (Daimon et al. 2003). Furthermore, inducible PLT5 over-expression sufficient for triggering ectopic cell proliferation in the shoot (Supplementary Fig. S8). We also observed overexpression of STM promoting ectopic shoot formation in *Arabidopsis* (Supplementary Fig. S8), as indicated in earlier studies (Scofield et al. 2008). To examine the consequence of down regulating STM, PLT5, and CUC2 in the *mea-*1^-/-^;*dme-*2^-/-^, we performed estradiol-inducible RNAi-mediated gene silencing. Upon estradiol-induced RNAi, *STM mRNA* levels reduced from 5.42-fold to 0.71-fold in *STM*-*dsRNAi*; *mea-*1^-/-^;*dme-* 2^-/-^ seedlings as compared to WT seedlings (Figure 5A). Similarly, *CUC2*-*dsRNAi* and *PLT5*-*dsRNAi* decreased *CUC2* and *PLT5 mRNA* levels from 2.52 and 3.54-fold to 0.93 and 0.6-fold, respectively (Figure 5A). Remarkably, ectopic shoot formation was significantly reduced upon reduction of *STM*, or *CUC2*, or *PLT5* transcripts in *mea-*1^-/-^;*dme-*2^-/-^ plants (Figure 5B). Estradiol-induced downregulation of *STM* in *mea-*1^-/-^;*dme-*2^-/-^ germinating seedlings reduced the frequency of ectopic shoot from 49% (N=4, n= 68/139, uninduced mock-treated seedlings; Supplementary Table 1) to 9% (N=4, n= 9/98, estradiol-induced seedlings), suggesting a partial rescue to WT phenotype (N=4, n= 89/98, Figure 5B, C). Similarly, RNAi-mediated knock-down of *CUC2* and *PLT5 mRNA* in *mea-*1^-/-^;*dme-*2^-/-^ germinating seedlings resulted in the reduction of ectopic shoots to 23% (N=3, n= 32/134) and 12% (N=4, n= 28/231), respectively (Figure 5B; Supplementary Fig. S9, Supplementary Table 1).

**Fig. 5.**
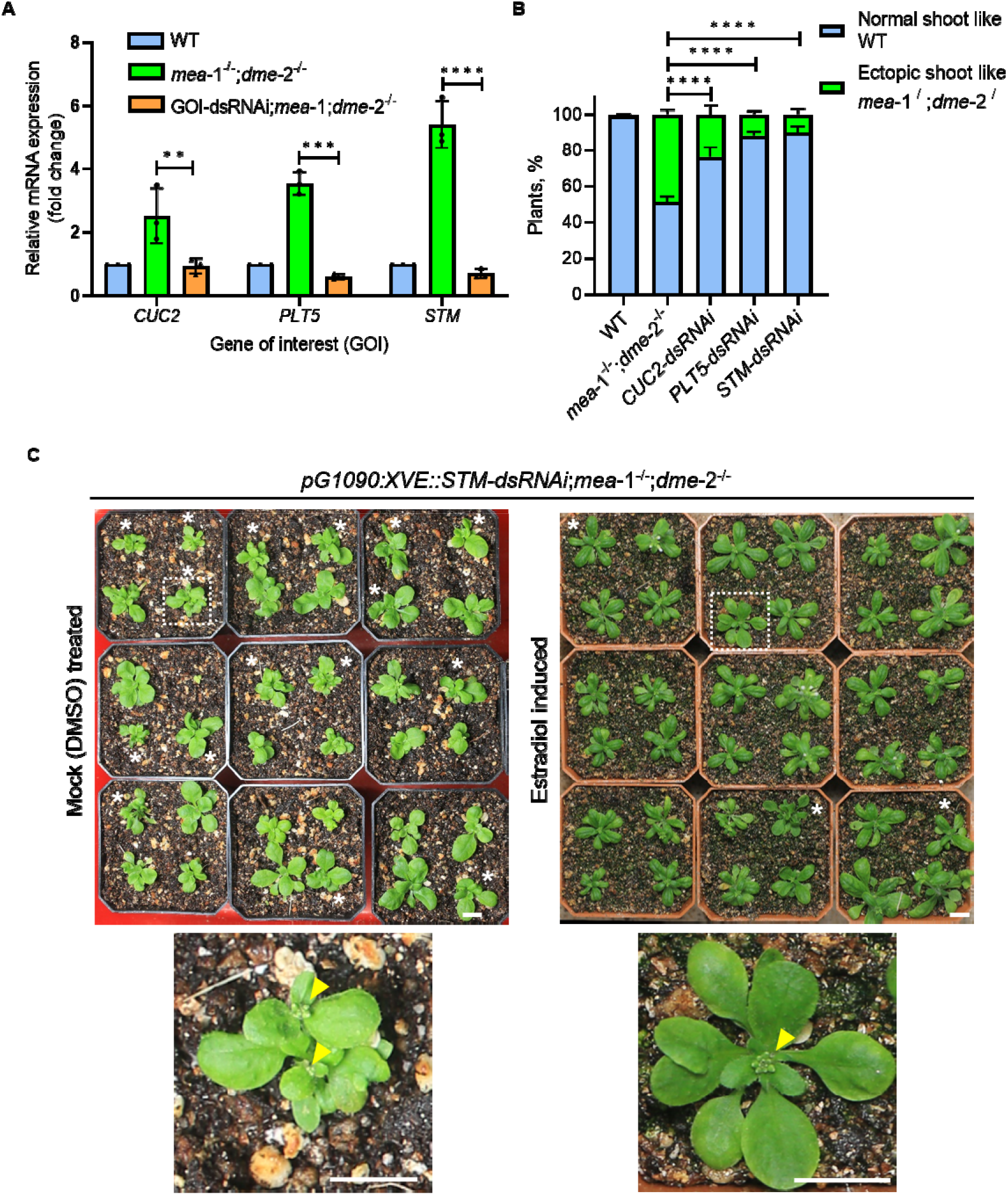
RNAi mediated downregulation of select target genes. (A) Relative abundance of *CUC2*, *PLT5*, and *STM* transcripts in respective RNAi lines. *CUC2*: WT-1±0, *mea-*1^-/-^;*dme-* 2^-/-^-2.52±0.87, *CUC2*-dsRNAi;*mea*-1-/-;*dme*-2-/--0.94±0.24, \**p*=0.038; *PLT5*: WT-1±0, *mea*-1-/-;*dme*-2-/--3.54±0.36, *PLT5*-dsRNAi;*mea*-1-/-;*dme*-2-/--0.60±0.08, \*\*\**p*=0.00048; *STM*: WT-1±0, *mea*-1-/-;*dme*-2-/--5.42±0.75, *STM*-dsRNAi;*mea*-1-/-;*dme*-2-/--0.71±0.15, \*\*\**p*=0.0008. Statistical test: Multiple unpaired *t*-tests (Holm-Sidak method), with alpha = 0.05. Error bar represents SD. **(B)** Quantification of shoot number in *CUC2*-, *PLT5*- and *STM*-dsRNAi lines as compared to WT and *mea*-1-/-;*dme*-2-/-. WT: n= 161 (Normal shoot= 161, ectopic shoot= 0); *mea*-1-/-;*dme*-2-/-: n= 139 (Normal shoot= 71, ectopic shoot= 68); *CUC2*-dsRNAi;*mea*-1-/-;*dme*-2-/-: n= 134 (Normal shoot= 102, ectopic shoot= 32, \*\*\*\**p*< 0.0001); *PLT5*-dsRNAi;*mea*-1-/-;*dme*-2-/-: n= 231 (Normal shoot= 203, ectopic shoot= 32, \*\*\*\**p*< 0.0001); *STM*-dsRNAi;*mea*-1-/-;*dme*-2-/-: n= 98 (Normal shoot= 89, ectopic shoot= 9, \*\*\*\**p*< 0.0001). Statistical analysis: two-way ANOVA (Sidak method), with alpha = 0.05. Error bar represents SEM. **(C)** Rescue of ectopic shoot in estradiol-induced *STM*-dsRNAi;*mea*-1-/-;*dme*-2-/-plants. Asterix(*) marks plants with ectopic shoots. White dotted box corresponds to the zoomed plants shown in the below panel. Yellow arrowhead points to the shoot apex. Scale bar represents 1cm.

Together, our results suggest that MEA and DME synergistically regulate shoot development by repressing the ectopic induction of shoot-promoting factors.

## Discussion

The histone methyltransferase, MEDEA, was identified as imprinted gene with the DNA glycosylase, DEMETER, facilitating its expression in the central cell of the female gametophyte. Both act synergistically to restrict the over-proliferation of the central cell in the absence of double fertilization (Gehring et al. 2006). Apart from peculiar seed abortion phenotype, hypomethylation by DME was discovered to aid H3K27me3 marks on the chromatin, linking it to PRC2 regulated pathway of gene expression (Moreno[Romero et al. 2016). Ovules inheriting the strong mutant alleles of both *mea* and *dme* abort seed development, making it impossible to recover homozygotes of either mutant. The maternal lethality of the *mea* and *dme* alleles makes it challenging to examine their roles beyond the gametophyte stage. However, recent studies have successfully implemented various genetic strategies to obtain viable, single mutant, *mea,* and *dme* homozygotes (Ravi et al. 2014; Pires et al. 2016; Kim et al. 2021; Williams et al. 2022). These single mutants have helped demonstrate the role of DME and MEA in various phases of sporophyte development, in addition to its prior established role in the gametophytes. For instance, MEA is required for proper embryonic patterning by regulating cell division via D-type cyclin 1.1 (CYCD1;1) (Simonini et al. 2021). MEA also facilitates *P*. *syringae* infection in *Arabidopsis* by downregulating RESISTANCE TO P. SYRINGAE2 (RPS2) (Roy et al. 2018). Like MEA, DME also has a sporophytic role in controlling cell division rate in sporophytic tissues like root hair and stomata, growth of inflorescence stem, and *de novo* shoot regeneration (Kim et al. 2021). DME, along with its somatic DNA demethylating counterparts, is also shown to regulate flowering time and inflorescence meristem activity (Kim et al. 2021; Williams et al. 2022). Like MEA, DME also restricts pathogen infection by repressing transposable elements controlling disease-related proteins in *Arabidopsis* (Schumann et al. 2019). In contrast to the documented single mutant phenotypes, *mea-*1^-/-^;*dme-*2^-/-^ plants show a unique presence of twin shoots arising from seed with a single embryo. This suggests the role of MEA and DME in controlling seed to seedling transition phase of sporophytic development.

The rescue of *mea-*1^-/-^;*dme-*2^-/-^ haploid sporophytes using CENH3-mediated genome elimination as a tool facilitated us to generate a population of *mea-*1^-/-^;*dme-*2^-/-^ DH plants which in turn shed light on the synergistic role of MEA and DME in suppressing the activation of ectopic shoot meristems during early sporophyte development. Shoot developmental processes like regeneration and floral transitions are known to be restricted by the repressive histone and DNA methylation marks (Li et al. 2011; Sun et al. 2014). However, only a few studies have reported the involvement of such epigenetic players during early sporophyte development (Bouyer et al. 2011; Gutzat et al. 2011). Seed to seedling transition is controlled by FIE, a subunit of the PRC2 complex (Bouyer et al. 2011). Here we propose that, during the sporophytic development, MEA, another subunit of the PRC2 complex, and DME restrict the formation of a single meristem at the apical pole by repressing shoot regulators. Consistent with this hypothesis, we show that an array of shoot regulators gets upregulated than WT levels in the *mea-*1^-/-^;*dme-*2^-/-^ mutants.

Restoration of shoot apex typical of wild type by down-regulation of a select few shoot-promoting factors in *mea-*1^-/-^;*dme-*2^-/-^ mutant strengthen this mechanistic framework. Epigenetic suppression of shoot-promoting factors such STM, PLT5, and CUC2 plausibly restrict the coordinated interactions between several fundamental cellular processes such as cell division and differentiation at the apical pole of the plant’s body axis during postembryonic development. It is important to note that MEA and DME do not share common targets. It is likely that MEA and DME mediated distinct regulatory inputs act synergistically to suppress ectopic shoot meristem formation.

In conclusion, we demonstrate an extended synergistic role for the gametophytic epigenetic regulators DME and MEA during the seed-to-seedling transition of sporophyte development. Our study provides the basic framework for understanding the role of gametophytic lethals in sporophytic tissue development. Further studies utilizing haploid induction for generating homozygous mutants may offer more insights into plant growth development.

## Materials and Methods

### Plant materials and growth conditions

Arabidopsis seeds were surface sterilized by fumigation with Chlorine (Cl_2_) gas (Lindsey et al. 2017). Fumigated seeds were then plated onto Murashige and Skoog (MS) growth medium with 0.8% agar and cold stratified at 4 [in dark for four days. After cold stratification, the plates were transferred to the growth chamber for germination. Seedlings and plants in pots were grown under white fluorescent lights (7000 lux at 20 cm) at 22 °C with a 16[h light/8 h dark cycle at 75% RH in a controlled growth cabinet (Percival Inc.). Heterozygous *MEA*/*mea*-1 (Grossniklaus et al. 1998) and *DME*/*dme*-2 (Choi et al. 2002) mutants were previously described. Following heterozygous seeds were obtained from Arabidopsis Biological Resource Center: *MEA*/*mea*-6 (CS6996), *MEA*/*mea*-7 (CS6997), *DME*/*dme*-6 (GK-252E03), and *DME*/*dme*-7 (SALK107538).

### Generation of maternal gametophytic lethal homozygotes

To generate *dme*-2 (AT5G04560) and *mea*-1 (AT1G02580) homozygous mutants, heterozygous *DME*/*dme*-2 and *MEA*/*mea*-1 plants were crossed as the male to haploid inducer GFP-*tailswap* plants as female (Ravi et al. 2014). The resultant F1 seeds were collected and germinated on Murashige and Skoog agar plates and later transferred to soil for further growth. Haploids were identified phenotypically and cytologically as described earlier (Ravi and Bondada 2016). All the haploids were PCR genotyped to identify the *DME* WT allele using MR146 and MR147 oligo combinations amplifying a 791 bp fragment. Similarly, the *dme*-2 mutant haploids were identified using MR147 and a left border-specific oligo MR148 producing 381 bp fragment. *MEA* allele was identified using oligonucleotides combinations MR8 and MR319 (722 bp), and the *mea*-1 allele using MR8 and a left border-specific oligo MR10 (312 bp). The DNA sequences for all oligos used in this study is given in Supplementary Table 5. The mutant haploids were screened for spontaneous mitotic/meiotic doubling events for obtaining doubled haploid seeds which gives rise to homozygous mutant diploid sporophytes. For generating homozygous *mea*-1;*dme*-2 double mutant in L*er* ecotype, *mea*-1 homozygous plants serving as female parent were crossed with *dme*-2 heterozygous plants as male parent. The resulting progeny PCR genotyped for both the *mea*-1 and *dme*-2 alleles, and the plants heterozygous plant both *mea*-1 and *dme*-2 alleles were identified and used as male parent in a cross with GFP-*tailswap* as female. *mea*-1;*dme*-2 haploid plants were identified using PCR-based genotyping. Spontaneous double haploid seeds from *mea*-1;*dme*-2 plants were collected to generate *mea*-1;*dme*-2 doubled haploid (diploid) sporophytes used in this study.

### Quantitative Real Time-PCR

For expression analysis of *DME* and *MEA* mRNA, we harvested shoot apices from 1, 3, 5, 7, and 10 dpg old WT seedlings and stored at -80 [for further use. Shoot apices from *mea-*1^-/-^;*dme-*2^-/-^ seedlings were used as control. Total RNA was isolated using TRIzol^TM^ reagent as per the manufacturer’s instructions. The extracted RNA was treated with NEB(New England Biolabs) DNase I (RNase free) (M0303) according to NEB guidelines to remove the genomic DNA contamination. One microgram of total RNA was used for complementary DNA (cDNA) synthesis by PrimeScript first strand cDNA synthesis kit (TAKARA-Bio, Cat# 6110) using an oligo(dT) and random hexamer as oligonucleotides. qPCRs were performed on Mastercycler BIORAD CFX96^TM^ using the oligonucleotides mentioned in Supplementary Table 5. *ACTIN2* (*ACT2*) served as a reference gene control. The reactions were carried out using TB Green® Premix Ex Taq™ II (TAKARA-Bio, Cat no. RR82WR) and incubated at 95 °C for 2 minutes, followed by 40 cycles of 95 °C for 15 s and 60 °C for 30 s. PCR specificity was checked by melting curve analysis. *ACT2* was used for the normalization for all the reactions. At least three independent biological samples were analyzed for each time point mentioned above, and qPCR reactions were set up with three technical replicates for each biological replica. mRNA abundance of target genes was analyzed (Supplementary Table 4) using the normalized 2^-ΔΔCt^ (cycle threshold) method or using absolute quantification utilizing standard curve (Livak and Schmittgen 2001).

### Microscopy and imaging

Bright-field images of seedlings were captured using Leica M205FA stereo microscope. Confocal laser scanning microscopic images were acquired using Zeiss LSM 880 and Leica TCS SP5 II confocal laser-scanning microscope. The confocal imaging was carried out on dissected embryos and 3 to 5 dpg old seedlings using Argon laser source at 40% power. The excitation wavelengths were as follows: 488 nmfor GFP and 514 nmfor YFP. Auto-fluorescence was captured at the wavelength of 633 nm. Images were captured with 40x (oil) or 63x (oil) objectives with 550-750 master gain, 100-200 range pinhole, and frame size of 1,024 by 1,024 pixels. Master gain and the pinhole adjustment varied with samples. The line averaging was adjusted to four and the digital zoom to 0.6. Z-stack images were taken in 5 µm or 1 µm intervals. The whole plant images are captured using a Canon 70D SLR camera. The images are edited in Adobe Photoshop CS4.

### ChIP followed by Next-Generation Sequencing

To identify the targets of MEA protein and differentiate histone methylation, we performed chromatin immunoprecipitation (ChIP) followed by next-generation sequencing as described previously (Yamaguchi et al. 2014; Salvi et al. 2020) on two independent biological replicates. 500 mg of micro dissected shoot apices, devoid of cotyledons and hypocotyl, from 2 to 3-day-old WT and *mea-*1^-/-^;*dme-*2^-/-^ seedlings were used for each sample. The samples were fixed in 1.8% formaldehyde in 1X PBS and vacuum infiltrated for 15 mins at room temperature to cross-link the tissue. After fixation, formaldehyde solution was removed and replaced by ice-cold 0.125 M glycine solution (in 1X PBS) followed by vacuum infiltration for 5 mins to quench the reaction. The glycine solution was filtered away and samples were washed twice with ice-cold 1X PBS. Samples were blotted on Kim wipe paper to remove excess 1X PBS. Samples were frozen and powdered in liquid nitrogen. The powdered samples were resuspended in nuclei extraction buffer (50 mM Tris-HCl pH 7.5, 150 mM NaCl, 1% Triton X-100, 0.1% sodium deoxycholate, 2.5 mM EDTA, 10% glycerol, freshly supplemented with cOmplete™, EDTA-free protease inhibitor cocktail (Cat# 04693132001) and 1 mM PMSF (Sigma). After brief incubation of 15 minutes, sample solutions were filtered using Falcon^®^ 40 µm cell strainer (Cat# 352340). Filtrate was centrifuged at 4°C to pellet the nuclei. The pellet was resuspended in an ice-cold nuclei lysis buffer (50 mM Tris-HCl, pH 8; 5 mM EDTA, pH 8; 0.5% SDS). The solution was sonicated for 8 cycles of 10 seconds with 20 seconds gap at 39% amplitude (Branson Sonifier, Marshall Scientific). 1.1% triton was added to the sonicated sample and centrifuged to isolate the supernatant. Preclearing of the sample was done by adding 20 μl of Dynabeads™ Protein G (Invitrogen, Cat# 10003D) and incubated at 4 [for 2h. An aliquot of the sample was stored as Input for the experiment. The sample was incubated with mouse monoclonal antibody to H3K27me3 (Cat# ab6002; Abcam) overnight at 4[, and Dynabeads™ Protein G was added to immunoprecipitate the DNA fragment for 5 hr at 4[. Sequential washes were performed in the beads using cold low salt wash buffer (150 mM NaCl, 0.2% SDS, 0.5% Triton X-100, 2 mM EDTA pH 8, 20 mM Tris-HCl pH 8), cold high salt wash buffer (500 mM NaCl, 0.2% SDS, 0.5% Triton X-100, 2 mM EDTA pH 8, 20 mM Tris-HCl pH 8), cold 250mM LiCl wash buffer (0.25 M LiCl, 0.5% NP-40, 0.5% sodium deoxycholate, 1 mM EDTA pH 8, 10 mM Tris-HCl pH 8) and 0.5X TE. All washes were done twice. Magnetic beads were collected and resuspended in TES (25 mM Tris-HCl pH 8, 5 mM EDTA pH 8, 0.5% SDS). 5 M NaCl was added to the sample and input control to reverse the cross-links overnight at 65[. DNA fragments were purified using a PCR purification kit (Cat #28106, Qiagen). Library constructions were performed, and validated using Qubit and Agilent TapeStation, followed by 150bp paired-end sequencing on Illumina NovaSeq 6000 by miBiome Therapeutics LLP, Mumbai, India.

The trimmed reads were aligned to reference genome TAIR10, using BWA 0.7.17-r1188 (Li 2013). The quality check of the alignments was done using Qualimap 2.2.2-dev (Okonechnikov et al. 2016). Duplicated reads, reads with low mapping quality, and mitochondrial and plastid reads were identified and removed with Samtools version 1.2 (Li et al. 2009). Enriched intervals between the IP sample and its relative Input were identified by MACS2 (version 2.0.10) program (Zhang et al. 2008) with the default setting. The big-Wig files were generated from the BAM files using Deeptools 3.5.1 (Ramírez et al. 2016). The coverage is calculated as the number of reads per bin and normalized to 1 x sequencing depth (reads per genome coverage, RPGC). The resulting bigWig files were visualized in Integrated Genome Viewer (https://igv.org/app/) (Robinson et al. 2011) to observe the protein binding on the chromatin. Peak annotation and GO and KEGG analysis for the target regions were done using ChIPseeker 1.32.1 (Yu et al. 2015) and ClusterProfiler 4.4.4 (Yu et al. 2012).

### Whole-genome bisulfite sequencing

To identify changes in DNA methylation, we performed bisulfite sequencing as described previously (Bondada et al. 2020). For this, we isolated genomic DNA from 500 mg of micro dissected shoot apices, devoid of cotyledons and hypocotyl, from 2 to 3-day-old WT and *mea-*1^-/-^;*dme-*2^-/-^ seedlings using NucleoSpin® Plant II (MACHEREY-NAGEL). 200 ng of DNA was subjected to fragmentation using ultrasonicator (Covaris) to get an average insert size of 250-300 bp. The fragments were end repaired, ligated to Enzymatic Methyl (EM)-seq adapters and excess adapter removed using AMPure XP purification beads. This was followed by two sets of enzymatic conversion steps using the Enzymatic Methyl-seq kit (NEB, Cat# E7120L) to differentiate cytosines from 5mC and 5hmC. The resulting converted sequences of the bisulfite-treated adapter ligated library were PCR enriched and clean-up of library products performed as instructed by the manufacturer. The cleaned libraries were quantitated on Qubit fluorometer and appropriate dilutions loaded on a HSD1000 screen tape to determine the size range of the fragments and the average library size.

Prepared library was 150bp paired-end sequenced on Illumina NovaSeq 6000 by miBiome Therapeutics LLP, Mumbai, India. Trimgalore 0.6.7 was used for the Illumina adapter removal. Each sample contained more than 40 million reads. FASTQC v0.11.9 was used for Illumina reads quality check. We used Bismark software utilizing Bowtie2 to align the clean reads to the TAIR10 reference genome, and more than 79% of the reads were uniquely mapped to the *Arabidopsis* genome in each sample (Supplementary Fig. S6). Bismark v0.24.0 was used to map the Illumina reads to the reference genome from TAIR10 (Krueger and Andrews 2011). Deduplication, extraction of methylation calls was also performed using Bismark v0.24.0. After removing duplicated reads, the number of methylated and unmethylated reads along with their trinucleotide context was generated for each cytosine on both the strands. The cytosines with overall coverage equal or more than five were considered for methylation calling. The cytosine with five or above-methylated reads or more than 40% of the reads were considered as methylated. Statistical analysis was performed using MethylKit 1.22.0 (Akalin et al. 2012). The differential methylation events were annotated for gene features using the ChIPseeker package (Yu et al. 2015).

### Plasmid construction, molecular cloning, and plant transformation

For cloning the gene and its regulatory sequences, the sequence of interest was PCR amplified from total genomic DNA isolated by CTAB method. For cDNA cloning, total RNA was isolated using TRIzol^TM^ reagent as per the manufacturer’s instructions. One microgram of DNase-free RNA was used for cDNA synthesis by PrimeScript 1st strand cDNA synthesis kit using an oligo(dT) and random hexamer as oligonucleotides. Polymerase chain reaction (PCR) was carried out using PrimeSTAR^®^ GXL DNA Polymerase (TAKARA-Bio, Cat# R050). The PCR conditions were specific to each sequence and amplified using 10 μM sequence-specific paired oligonucleotides (see Supplementary Table 5) using Applied Biosystems™ ProFlex™ 3 x 32-well PCR System. All entry clones were propagated in DH5α strain of *E*.*coli* using a single Gateway cloning system. To knock down the expression of *STM*, 272 bp exon sequence was amplified from *cDNA* using region-specific oligonucleotides and cloned in the p2R3E-As-OScT vector (sense) and p221Z-S-Intr vector (antisense) using *Bam*HI and *Xho*I restriction digestion and ligation. All entry clones were propagated in the DH5 [strain of *E*.*coli* using a single gateway cloning system (Invitrogen, Cat# 12538120). The sense and antisense for STM were cloned under *pG1090* inducible promoter (Zuo et al. 2000) and subsequently in *pCAMBIA-based R4R3* destination vector. Similarly, the *PLT5* (299 bp) and *CUC2* (Varapparambath et al. 2022) knockdown constructs were cloned under *pG1090* inducible promoter to silence the transcript level of *PLT5* and *CUC2*, respectively. To generate *pDME*::*3XVENUS-NLS*:*3AT*, 5000bp upstream regulatory sequence from start codon of *DME* is tagged with *3XVENUS* with nuclear localization signal (NLS) and *3AT* terminator. Similarly, 5142bp upstream regulatory sequence from start codon of *MEA* was tagged with *3XVENUS* with nuclear localization signal (NLS) to generate *pMEA*::*3XVENUS-NLS*:*3AT*. The transcriptional reporter for *WUS* and CUC2 protein fusion were described previously (Radhakrishnan et al. 2020; Varapparambath et al. 2022). These constructs were introduced into *Agrobacterium tumefaciens* strain C58 by electroporation. Stable transgenic plants were generated by floral-dip method (Clough and Bent 1998).

### Estradiol treatment for inducible gene knock-down

Inducible RNAi for *STM*, *PLT5*, and *CUC2* genes was carried out, as mentioned earlier (Varapparambath et al. 2022). MS agar medium supplemented with 5μM β-estradiol was used to induce conditional gene silencing using *pG1090*:*XVE* artificial promoter throughout seed germination. Estradiol-treated and mock (DMSO) treated seedlings were scored for ectopic shoot emergence at 15 dpg.

### Statistical analysis

The details of the statistical analysis followed in this study are described in the respective figure legends (number of samples (n), mean± S.E.M). The statistical significance levels were defined as \**p* < 0.05, \*\**p* < 0.01, \*\*\**p* < 0.001, \*\*\*\**p* < 0.0001. For mRNA quantitative analysis, statistical significance was determined using the Multiple unpaired *t*-tests (Holm-Sidak method), with alpha = 0.05. For RNAi phenotype quantitative analysis, statistical significance was determined using two-way ANOVA (Sidak method), with alpha = 0.05. All the statistical tests and analyses were performed using GraphPad PRISM (v9.5.1) statistical software.

## Data Availability

The ChIP-sequencing and whole-genome bisulfite sequencing data generated and analyzed during the current study are available from NCBI under the bio project accession PRJNA972183 consisting of eight SRA accession identifiers: SRR24547737, SRR24547736, SRR24547733, SRR24547734, SRR24547735, SRR24547732, SRR25392706 and SRR25392707. All other source data are included in the article as Supplementary Table 2 and 3.

## Acknowledgments

M.P.R. thanks to the IISER Thiruvananthapuram for providing a Ph.D. research fellowship. S.S. is a recipient of the Department of Science and Technology (DST) - INSPIRE scholarship for higher education (SHE). R.B. is the recipient of a Junior and Senior Research Fellowship (J.R.F. and S.R.F.) for a Ph.D. funded by the University Grants Commission (U.G.C.), Govt. of India. A.P.S. thanks to the Council of Scientific and Industrial Research (CSIR) for the Ph.D. fellowship. K.P. acknowledges grants from the Department of Biotechnology (D.B.T.), Government of India (<u>BT/PR41931/BRB/10/1964/2021</u>), the Science and Engineering Research Board-Scientific and Useful Profound Research Advancement (SERB-SUPRA), Government of India (<u>S.P.R./2021/000109</u>). R.M. acknowledges financial support from Ramalingaswami’s re-entry fellowship awarded by the Department of Biotechnology (D.B.T., Govt. of India) and the Dupont Young Professor grant, Dupont U.S.A. We thank Dr. Ajith V.P for help in ChIP-Seq data analysis and Dr. Nishant K.T and Sameer Joshi for discussion and performing chromosome dosage analysis. We thank the Indian Institute of Science Education and Research (IISER)-Thiruvananthapuram for intramural financial and infrastructural support.

## Author contributions

R.M. conceptualized, designed, and supervised the study. M.P.R. contributed to certain aspects of the study design and carried out most of the experiments. S.S. generated and performed the phenotypic assessment of RNAi lines along M.P.R. R.B. helped generate double mutant lines. A.P.S. performed cloning experiments for producing reporter lines. K.P. designed, supervised, and critically analyzed the study. M.P.R., K.P., and R.M. wrote the manuscript. R.M. and K.P. acquired funds for the study. All the authors read and approved the draft.

## References

Aichinger E, Villar CBR, di Mambro R, et al (2011) The CHD3 chromatin remodeler PICKLE and polycomb group proteins antagonistically regulate meristem activity in the Arabidopsis root. Plant Cell 23:1047–1060. 10.1105/tpc.111.083352

Aida M, Ishida T, Tasaka M (1999) Shoot apical meristem and cotyledon formation during Arabidopsis embryogenesis: Interaction among the CUP-SHAPED COTYLEDON and SHOOT MERISTEMLESS genes. Development 126:1563–1570. 10.1242/dev.126.8.1563

Aida M, Vernoux T, Furutani M, et al (2002) Roles of PIN-FORMED1 and MONOPTEROS in pattern formation of the apical region of the Arabidopsis embryo. Development 129:3965–3974. 10.1242/dev.129.17.3965

Akalin A, Kormaksson M, Li S, et al (2012) methylKit: a comprehensive R package for the analysis of genome-wide DNA methylation profiles. Genome Biol 13:R87. 10.1186/gb-2012-13-10-r87

Baubec T, Finke A, Mittelsten Scheid O, Pecinka A (2014) Meristem[specific expression of epigenetic regulators safeguards transposon silencing in Arabidopsis. EMBO Rep 15:446–452. 10.1002/embr.201337915

Bondada R, Somasundaram S, Marimuthu MP, et al (2020) Natural epialleles of Arabidopsis SUPERMAN display superwoman phenotypes. Commun Biol 3:1–13. 10.1038/s42003-020-01525-9

Borg M, Jacob Y, Susaki D, et al (2020) Targeted reprogramming of H3K27me3 resets epigenetic memory in plant paternal chromatin. Nat Cell Biol 22:621–629. 10.1038/s41556-020-0515-y

Bouyer D, Kramdi A, Kassam M, et al (2017) DNA methylation dynamics during early plant life. Genome Biol 18:1–12. 10.1186/s13059-017-1313-0

Bouyer D, Roudier F, Heese M, et al (2011) Polycomb repressive complex 2 controls the embryo-to-seedling phase transition. PLoS Genet 7:. 10.1371/journal.pgen.1002014

Calarco JP, Borges F, Donoghue MTA, et al (2012) Reprogramming of DNA methylation in pollen guides epigenetic inheritance via small RNA. Cell 151:194–205. 10.1016/j.cell.2012.09.001

Chen D, Molitor A, Liu C, Shen WH (2010) The arabidopsis PRC1-like ring-finger proteins are necessary for repression of embryonic traits during vegetative growth. Cell Res 20:1332–1344. 10.1038/cr.2010.151

Chen D, Molitor AM, Xu L, Shen W-H (2016) Arabidopsis PRC1 core component AtRING1 regulates stem cell-determining carpel development mainly through repression of class I KNOX genes. BMC Biol 14:112. 10.1186/s12915-016-0336-4

Choi Y, Gehring M, Johnson L, et al (2002) DEMETER, a DNA glycosylase domain protein, is required for endosperm gene imprinting and seed viability in Arabidopsis. Cell 110:33–42. 10.1016/S0092-8674(02)00807-3

Clough SJ, Bent AF (1998) Floral dip: a simplified method for Agrobacterium-mediated transformation of Arabidopsis thaliana. Plant J 16:735–743. 10.1046/j.1365-313x.1998.00343.x

Cui X, Lu F, Qiu Q, et al (2016) REF6 recognizes a specific DNA sequence to demethylate H3K27me3 and regulate organ boundary formation in Arabidopsis. Nat Genet 48:694–699. 10.1038/ng.3556

Daimon Y, Takabe K, Tasaka M (2003) The CUP-SHAPED COTYLEDON Genes Promote Adventitious Shoot Formation on Calli. Plant Cell Physiol 44:113–121. 10.1093/pcp/pcg038

Dasgupta P, Chaudhuri S (2019) Analysis of DNA methylation profile in plants by chop-PCR. In: Methods in Molecular Biology. pp 79–90

De Lucia F, Crevillen P, Jones AME, et al (2008) A PHD-polycomb repressive complex 2 triggers the epigenetic silencing of FLC during vernalization. Proc Natl Acad Sci U S A 105:16831–16836. 10.1073/pnas.0808687105

Du J, Johnson LM, Jacobsen SE, Patel DJ (2015) DNA methylation pathways and their crosstalk with histone methylation. Nat Rev Mol Cell Biol 16:519–532. 10.1038/nrm4043

Feng S, Jacobsen SE, Reik W (2010) Epigenetic Reprogramming in Plant and Animal Development. Science 330:622–627. 10.1126/science.1190614

Figueiredo DD, Batista RA, Roszak PJ, Köhler C (2015) Auxin production couples endosperm development to fertilization. Nat Plants 1:1–6. 10.1038/nplants.2015.184

Gehring M, Huh JH, Hsieh T-FF, et al (2006) DEMETER DNA Glycosylase Establishes MEDEA Polycomb Gene Self-Imprinting by Allele-Specific Demethylation. Cell 124:495–506. 10.1016/j.cell.2005.12.034.

Grossniklaus U, Vielle-Calzada JP, Hoeppner MA, Gagliano WB (1998) Maternal control of embryogenesis by MEDEA, a Polycomb group gene in Arabidopsis. Science 280:446–450. 10.1126/science.280.5362.446

Guitton A-E, Page DR, Chambrier P, et al (2004) Identification of new members of Fertilisation Independent Seed Polycomb Group pathway involved in the control of seed development in Arabidopsis thaliana. Development 131:2971–81. 10.1242/dev.01168

Gutzat R, Borghi L, Fütterer J, et al (2011) RETINOBLASTOMA-RELATED PROTEIN controls the transition to autotrophic plant development. Development 138:2977–86. 10.1242/dev.060830

Gutzat R, Rembart K, Nussbaumer T, et al (2020) *Arabidopsis* shoot stem cells display dynamic transcription and DNA methylation patterns. EMBO J 39:e103667. 10.15252/embj.2019103667

Heisler MG, Ohno C, Das P, et al (2005) Patterns of auxin transport and gene expression during primordium development revealed by live imaging of the Arabidopsis inflorescence meristem. Curr Biol 15:1899–1911. 10.1016/j.cub.2005.09.052

Higo A, Saihara N, Miura F, et al (2020) DNA methylation is reconfigured at the onset of reproduction in rice shoot apical meristem. Nat Commun 11:1–12. 10.1038/s41467-020-17963-2

Hofhuis H, Laskowski M, Du Y, et al (2013) Phyllotaxis and rhizotaxis in Arabidopsis are modified by three plethora transcription factors. Curr Biol 23:956–962. 10.1016/j.cub.2013.04.048

Ibarra CA, Feng X, Schoft VK, et al (2012) Active DNA demethylation in plant companion cells reinforces transposon methylation in gametes. Science 337:1360–1364. 10.1126/science.1224839

Iwasaki M, Paszkowski J (2014) Epigenetic memory in plants. EMBO J 33:1987–1998. 10.15252/embj.201488883

Jullien PE, Susaki D, Yelagandula R, et al (2012) DNA Methylation Dynamics during Sexual Reproduction in Arabidopsis thaliana. Curr Biol 22:1825–1830. 10.1016/j.cub.2012.07.061

Kareem A, Durgaprasad K, Sugimoto K, et al (2015) PLETHORA genes control regeneration by a two-step mechanism. Curr Biol 25:1017–1030. 10.1016/j.cub.2015.02.022

Khouider S, Borges F, LeBlanc C, et al (2021) Male fertility in Arabidopsis requires active DNA demethylation of genes that control pollen tube function. Nat Commun 12:1–10. 10.1038/s41467-020-20606-1

Kim M, Ohr H, Lee JW, et al (2008) Temporal and Spatial Downregulation of Arabidopsis MET1 Activity Results in Global DNA Hypomethylation and Developmental Defects. Mol Cells 26:611–615

Kim S, Park J-SS, Lee J, et al (2021) The DME demethylase regulates sporophyte gene expression, cell proliferation, differentiation, and meristem resurrection. Proc Natl Acad Sci U S A 118:. 10.1073/pnas.2026806118

Köhler C, Hennig L, Spillane C, et al (2003) The Polycomb-group protein MEDEA regulates seed development by controlling expression of the MADS-box gene PHERES1. Genes Dev 17:1540–1553. 10.1101/gad.257403.431

Krueger F, Andrews SR (2011) Bismark: a flexible aligner and methylation caller for Bisulfite-Seq applications. Bioinformatics 27:1571–1572. 10.1093/bioinformatics/btr167

Kwon CS, Hibara KI, Pfluger J, et al (2006) A role for chromatin remodeling in regulation of CUC gene expression in the Arabidopsis cotyledon boundary. Development 133:3223–3230. 10.1242/dev.02508

Lafos M, Kroll P, Hohenstatt ML, et al (2011) Dynamic Regulation of H3K27 Trimethylation during Arabidopsis Differentiation. PLOS Genet 7:e1002040. 10.1371/journal.pgen.1002040

Lee K, Seo PJ (2018) Dynamic Epigenetic Changes during Plant Regeneration. Trends Plant Sci 23:235–247. 10.1016/j.tplants.2017.11.009

Li H (2013) Aligning sequence reads, clone sequences and assembly contigs with BWAMEM. ArXiv Prepr 1303:3997

Li H, Handsaker B, Wysoker A, et al (2009) The Sequence Alignment/Map format and SAMtools. Bioinformatics 25:2078–2079. 10.1093/bioinformatics/btp352

Li ZH, Lu X, Gao Y, et al (2011) Polyploidization and epigenetics. Chin Sci Bull 56:245–252. 10.1007/s11434-010-4290-1

Lindsey BE, Rivero L, Calhoun CS, et al (2017) Standardized method for high-throughput sterilization of Arabidopsis seeds. J Vis Exp 2017:1–7. 10.3791/56587

Livak KJ, Schmittgen TD (2001) Analysis of Relative Gene Expression Data Using Real-Time Quantitative PCR and the 2−ΔΔCT Method. Methods 25:402–408. 10.1006/meth.2001.1262

Lodha M, Marco CF, Timmermans MCP (2013) The ASYMMETRIC LEAVES complex maintains repression of KNOX homeobox genes via direct recruitment of Polycombrepressive complex2. Genes Dev 27:596–601. 10.1101/gad.211425.112

Luo M, Bilodeau P, Dennis ES, et al (2000) Expression and parent-of-origin effects for FIS2, MEA, and FIE in the endosperm and embryo of developing Arabidopsis seeds. Proc Natl Acad Sci U S A 97:10637–10642. 10.1073/pnas.170292997

Mayer KFX, Schoof H, Haecker A, et al (1998) Role of WUSCHEL in Regulating Stem Cell Fate in the Arabidopsis Shoot Meristem. Cell 95:805–815. 10.1111/j.1365-2818.1977.tb01111.x

Moreno[Romero J, Jiang H, Santos[González J, Köhler C (2016) Parental epigenetic asymmetry of PRC 2[mediated histone modifications in the Arabidopsis endosperm. EMBO J 35:1298–1311. 10.15252/embj.201593534

Okonechnikov K, Conesa A, García-Alcalde F (2016) Qualimap 2: Advanced multi-sample quality control for high-throughput sequencing data. Bioinformatics 32:292–294. 10.1093/bioinformatics/btv566

Park JS, Frost JM, Park K, et al (2017) Control of DEMETER DNA demethylase gene transcription in male and female gamete companion cells in Arabidopsis thaliana. Proc Natl Acad Sci U S A 114:2078–2083. 10.1073/pnas.1620592114

Park K, Lee S, Yoo H, Choi Y (2020) DEMETER-mediated DNA Demethylation in Gamete Companion Cells and the Endosperm, and its Possible Role in Embryo Development in Arabidopsis. J Plant Biol 63:321–329. 10.1007/s12374-020-09258-2

Pires ND, Bemer M, Müller LM, et al (2016) Quantitative Genetics Identifies Cryptic Genetic Variation Involved in the Paternal Regulation of Seed Development. PLOS Genet 12:e1005806. 10.1371/journal.pgen.1005806

Prasad K, Grigg SP, Barkoulas M, et al (2011) Arabidopsis PLETHORA transcription factors control phyllotaxis. Curr Biol 21:1123–1128. 10.1016/j.cub.2011.05.009

Qiu Q, Mei H, Deng X, et al (2019) DNA methylation repels targeting of Arabidopsis REF6. Nat Commun 10:1–9. 10.1038/s41467-019-10026-1

Radhakrishnan D, Shanmukhan AP, Kareem A, et al (2020) A coherent feed-forward loop drives vascular regeneration in damaged aerial organs of plants growing in a normal developmental context. Development 147:dev185710. 10.1242/dev.185710

Ramírez F, Ryan DP, Grüning B, et al (2016) deepTools2: a next generation web server for deep-sequencing data analysis. Nucleic Acids Res 44:W160–W165. 10.1093/NAR/GKW257

Ravi M, Bondada R (2016) Genome Elimination by Tailswap CenH3: In Vivo Haploid Production in Arabidopsis thaliana. In: Chromosome and Genomic Engineering in Plants: Methods and Protocols, Methods in Molecular Biology

Ravi M, Chan SWL (2010) Haploid plants produced by centromere-mediated genome elimination. Nature 464:615–618. 10.1038/nature08842

Ravi M, Marimuthu MPA, Tan EH, et al (2014) A haploid genetics toolbox for Arabidopsis thaliana. Nat Commun 5:1–8. 10.1038/ncomms6334

Robinson JT, Thorvaldsdóttir H, Winckler W, et al (2011) Integrative genomics viewer. Nat Biotechnol 29:24–26. 10.1038/nbt.1754

Roszak PJ, Köhler C (2011) Polycomb group proteins are required to couple seed coat initiation to fertilization. Proc Natl Acad Sci U S A 108:20826–20831. 10.1073/pnas.1117111108

Roy S, Gupta P, Rajabhoj MP, et al (2018) The polycomb-group repressor MEDEA attenuates pathogen defense. Plant Physiol 177:1728–1742. 10.1104/pp.17.01579

Salvi E, Rutten JP, Di Mambro R, et al (2020) A Self-Organized PLT/Auxin/ARR-B Network Controls the Dynamics of Root Zonation Development in Arabidopsis thaliana. Dev Cell 53:431–443.e23. 10.1016/j.devcel.2020.04.004

Schmid M, Davison TS, Henz SR, et al (2005) A gene expression map of Arabidopsis thaliana development. Nat Genet 37:501–506. 10.1038/ng1543

Schoft VK, Chumak N, Choi Y, et al (2011) Function of the DEMETER DNA glycosylase in the Arabidopsis thaliana male gametophyte. Proc Natl Acad Sci USA 108:8042–7. 10.1073/pnas.1105117108

Schoof H, Lenhard M, Haecker A, et al (2000) The stem cell population of Arabidopsis shoot meristems is maintained by a regulatory loop between the CLAVATA and WUSCHEL genes. Cell 100:635–644. 10.1016/S0092-8674(00)80700-X

Schumann U, Lee JM, Smith NA, et al (2019) DEMETER plays a role in DNA demethylation and disease response in somatic tissues of Arabidopsis. Epigenetics 14:1074–1087. 10.1080/15592294.2019.1631113

Scofield S, Dewitte W, Murray JA (2008) A model for Arabidopsis class-1 KNOX gene function. Plant Signal Behav 3:257–259. 10.4161/psb.3.4.5194

Shen W-H, Xu L (2009) Chromatin Remodeling in Stem Cell Maintenance in Arabidopsis thaliana. Mol Plant 2:600–609. 10.1093/mp/ssp022

Simonini S, Bemer M, Bencivenga S, et al (2021) The Polycomb group protein MEDEA controls cell proliferation and embryonic patterning in Arabidopsis. Dev Cell 56:1945–1960.e7. 10.1016/j.devcel.2021.06.004

Slotkin RK, Vaughn M, Borges F, et al (2009) Epigenetic Reprogramming and Small RNA Silencing of Transposable Elements in Pollen. Cell 136:461–472. 10.1016/j.cell.2008.12.038

Spillane C, Schmid KJ, Laoueillé-Duprat S, et al (2007) Positive darwinian selection at the imprinted MEDEA locus in plants. Nature 448:349–352. 10.1038/nature05984

Spinelli SV, Martin AP, Viola IL, et al (2011) A Mechanistic Link between *STM* and *CUC1* during Arabidopsis Development. Plant Physiol 156:1894–1904. 10.1104/pp.111.177709

Stahl Y, Simon R (2010) Plant primary meristems: shared functions and regulatory mechanisms. Curr Opin Plant Biol 13:53–58. 10.1016/j.pbi.2009.09.008

Sun B, Looi LS, Guo S, et al (2014) Timing mechanism dependent on cell division is invoked by Polycomb eviction in plant stem cells. Science 343:. 10.1126/science.1248559

Tan EH, Henry IM, Ravi M, et al (2015) Catastrophic chromosomal restructuring during genome elimination in plants. eLife 4:1–16. 10.7554/eLife.06516

Ten Hove CA, Lu KJ, Weijers D (2015) Building a plant: Cell fate specification in the early arabidopsis embryo. Dev Camb 142:420–430. 10.1242/dev.111500

Varapparambath V, Mathew MM, Shanmukhan AP, et al (2022) Mechanical conflict caused by a cell-wall-loosening enzyme activates de novo shoot regeneration. Dev Cell 57:2063–2080. 10.1016/j.devcel.2022.07.017

Vielle-Calzada JP, Thomas J, Spillane C, et al (1999) Maintenance of genomic imprinting at the Arabidopsis medea locus requires zygotic DDM1 activity. Genes Dev 13:2971– 2982. 10.1101/gad.13.22.2971

Williams BP, Bechen LL, Pohlmann DA, Gehring M (2022) Somatic DNA demethylation generates tissue-specific methylation states and impacts flowering time. Plant Cell 34:1189–1206. 10.1093/plcell/koab319

Williams BP, Gehring M (2020) Principles of Epigenetic Homeostasis Shared Between Flowering Plants and Mammals. Trends Genet 36:751–763. 10.1016/j.tig.2020.06.019

Xiao J, Jin R, Yu X, et al (2017) Cis and trans determinants of epigenetic silencing by Polycomb repressive complex 2 in Arabidopsis. Nat Genet 49:1546–1552. 10.1038/ng.3937

Xiao J, Lee U-S, Wagner D (2016) Tug of war: adding and removing histone lysine methylation in Arabidopsis. Curr Opin Plant Biol 34:41–53. 10.1016/j.pbi.2016.08.002

Xiao J, Wagner D (2015) Polycomb repression in the regulation of growth and development in Arabidopsis. Curr Opin Plant Biol 23:15–24. 10.1016/j.pbi.2014.10.003

Xiao W, Gehring M, Choi Y, et al (2003) Imprinting of the MEA polycomb gene is controlled by antagonism between MET1 methyltransferase and DME glycosylase. Dev Cell 5:891–901. 10.1016/S1534-5807(03)00361-7

Xu L, Shen WH (2008) Polycomb Silencing of KNOX Genes Confines Shoot Stem Cell Niches in Arabidopsis. Curr Biol 18:1966–1971. 10.1016/j.cub.2008.11.019

Yamaguchi N, Winter CM, Wu M-F, et al (2014) PROTOCOL: Chromatin Immunoprecipitation from Arabidopsis Tissues. In: The Arabidopsis Book. p e0170

Yoshida N, Yanai Y, Chen L, et al (2001) EMBRYONIC FLOWER2, a novel Polycomb group protein homolog, mediates shoot development and flowering in Arabidopsis. Plant Cell 13:2471–2481. 10.1105/tpc.13.11.2471

You Y, Sawikowska A, Neumann M, et al (2017) Temporal dynamics of gene expression and histone marks at the Arabidopsis shoot meristem during flowering. Nat Commun 8:15120. 10.1038/ncomms15120

Yu G, Wang LG, Han Y, He QY (2012) ClusterProfiler: An R package for comparing biological themes among gene clusters. OMICS J Integr Biol 16:284–287. 10.1089/omi.2011.0118

Yu G, Wang LG, He QY (2015) ChIP seeker: An R/Bioconductor package for ChIP peak annotation, comparison and visualization. Bioinformatics 31:2382–2383. 10.1093/bioinformatics/btv145

Zhang H, Tang K, Wang B, et al (2014) Protocol: A beginner’s guide to the analysis of RNA-directed DNA methylation in plants. Plant Methods 10:1–9. 10.1186/1746-4811-10-18

Zhang S, Wang D, Zhang H, et al (2018) FERTILIZATION-INDEPENDENT SEED-Polycomb Repressive Complex 2 Plays a Dual Role in Regulating Type i MADS-Box Genes in Early Endosperm Development. Plant Physiol 177:285–299. 10.1104/PP.17.00534

Zhang X, Clarenz O, Cokus S, et al (2007) Whole-Genome Analysis of Histone H3 Lysine 27 Trimethylation in Arabidopsis. PLoS Biol 5:e129. 10.1371/journal.pbio.0050129

Zhang Y, Liu T, Meyer CA, et al (2008) Model-based Analysis of ChIP-Seq (MACS). Genome Biol 9:R137. 10.1186/gb-2008-9-9-r137

Zuo J, Niu QW, Chua NH (2000) An estrogen receptor-based transactivator XVE mediates highly inducible gene expression in transgenic plants. Plant J 24:265–273. 10.1046/j.1365-313X.2000.00868.x

