## Supplementary Data for "Gametophytic epigenetic regulators MEDEA and DEMETER synergistically suppress ectopic shoot formation in Arabidopsis"

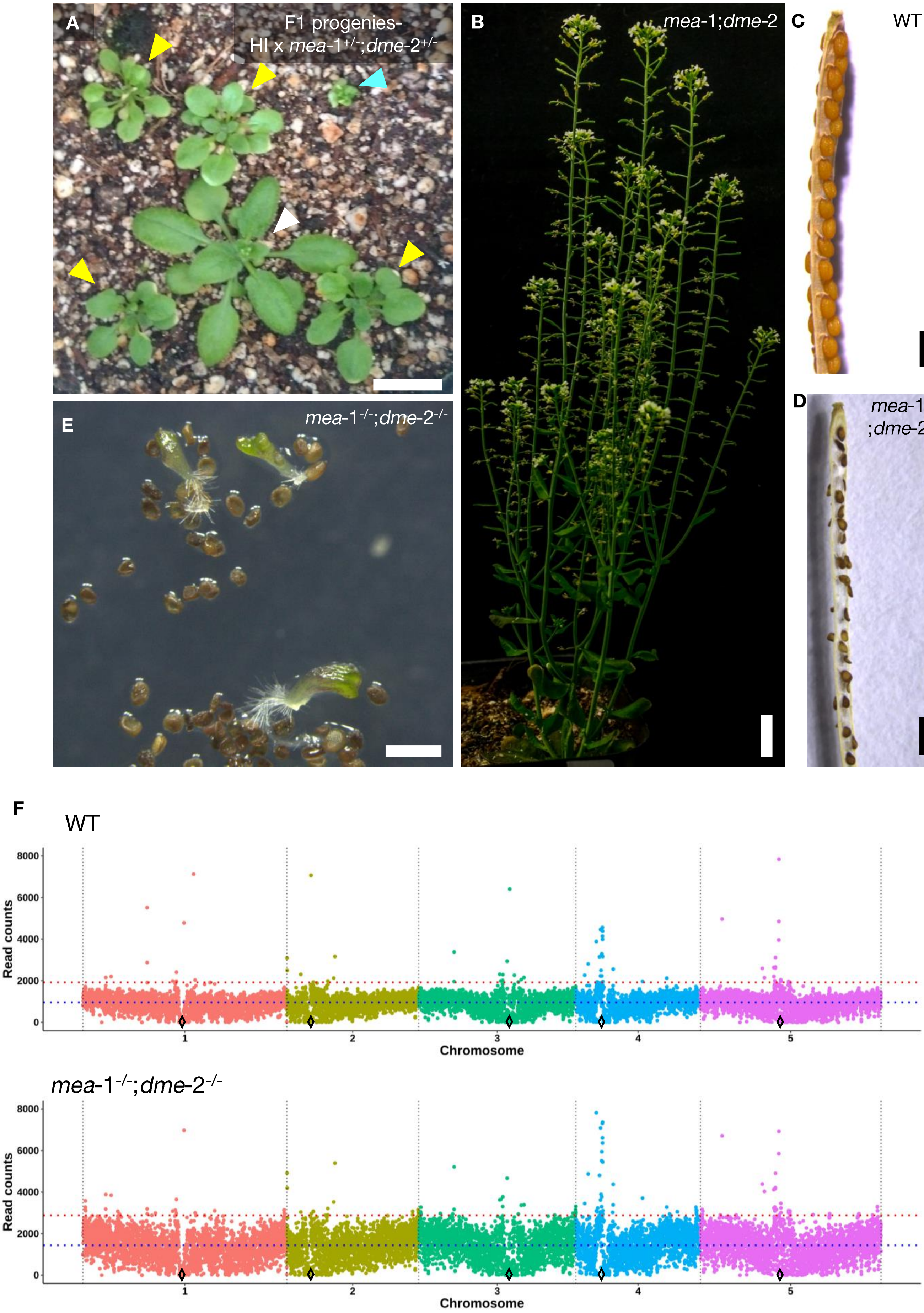
A
F1 progenies-
HI x mea-1+/-;dme-2+/-
B
mea-1;dme-2
C
WT
D
mea-1-/-
;dme-2-/-
E
mea-1-/-;dme-2-/-
F
WT
Read counts
8000
6000
4000
2000
0
1
2
3
4
5
Chromosome
mea-1-/-;dme-2-/-
Read counts
8000
6000
4000
2000
0
1
2
3
4
5
Chromosome

**Supplementary Fig. S1 Phenotypes of HI x *mea*-1$^{+/-}$;*dme*-2$^{+/-}$ F1 and *mea*-1$^{-/-}$;*dme*-2$^{-/-}$ DH progeny along with its chromosome dosage analysis**

**(A)** F1 progeny obtained from HI x *mea*-1$^{+/-}$;*dme*-2$^{+/-}$ cross. Haploid plants (yellow arrow) exhibit reduced size as compare to diploid *mea*-1$^{+/-}$;*dme*-2$^{+/-}$ (L*er*) (white arrow) and are dissimilar from aneuploid plants (blue arrow), which display distinct phenotypes

**(B)** *mea*-1;*dme*-2 haploid plant

**(C)** WT silique displaying viable seeds

**(D)** doubled haploid *mea*-1$^{-/-}$;*dme*-2$^{-/-}$ silique displaying non viable (shriveled, dark brown) seeds

**(E)** *mea*-1$^{-/-}$;*dme*-2$^{-/-}$ seeds exhibiting low germination along with seedlings depicting abnormal phenotype. The scale bar represents **(A-B)** 1 cm **(C-E)** 1 mm

**(F)** Chromosome dosage analysis of *mea*-1$^{-/-}$;*dme*-2$^{-/-}$ seedlings as compared to diploid WT showing no numerical or structural aneuploidy, based on 100 kb bin size. Black diamond indicate relative centromere positions.

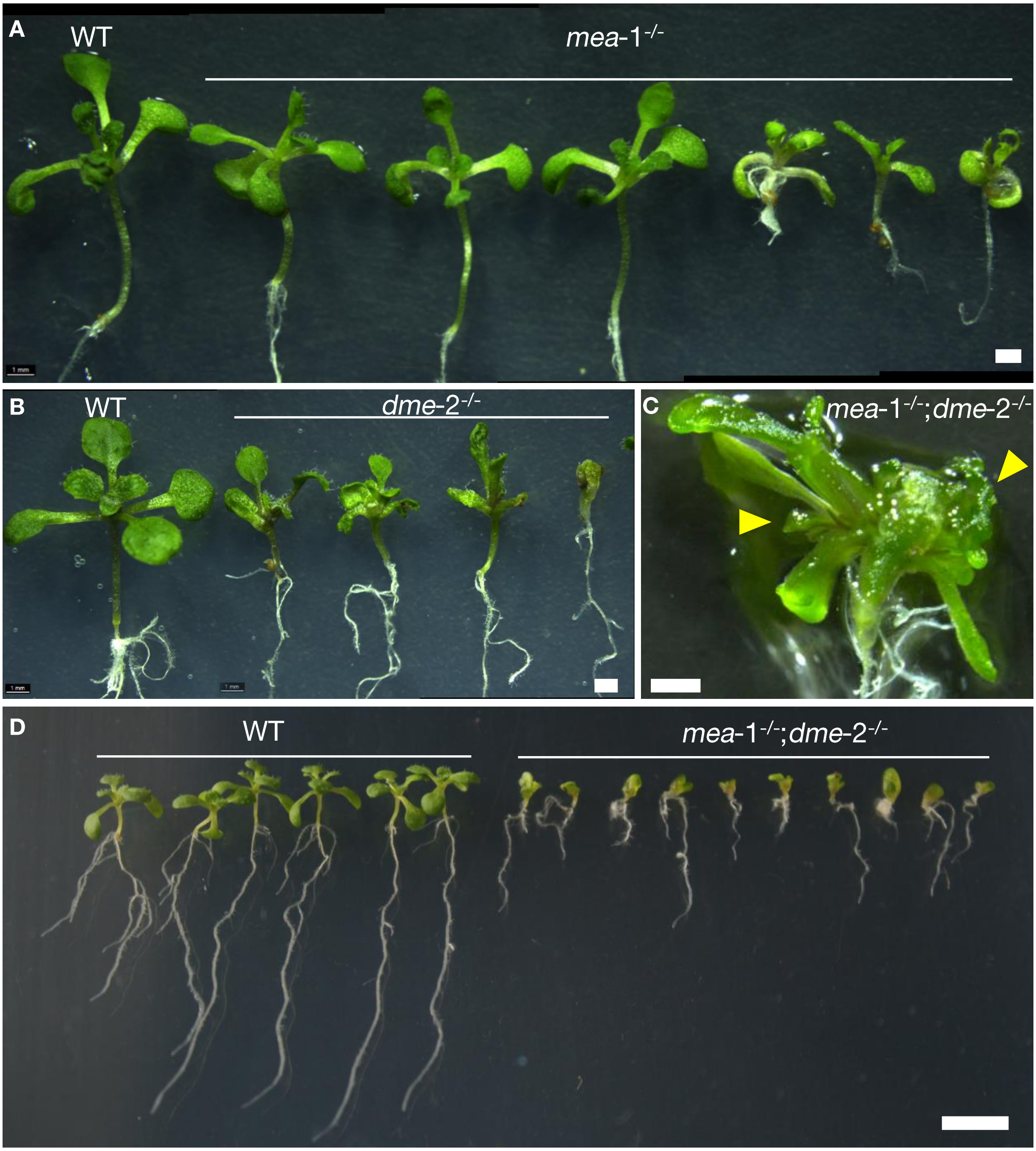


**Supplementary Fig. S2 Phenotypes of single mutant *mea*-1$^{-/-}$, *dme*-2$^{-/-}$ and double mutant *mea*-1$^{-/-}$;*dme*-2$^{-/-}$ seedlings**

**(A)** *mea*-1$^{-/-}$ seedlings as compared to WT at 6-leaf (10 dpg) stage

**(B)** *dme*-2$^{-/-}$ seedlings as compared to WT at 6-leaf (10 dpg) stage

**(C)** *mea*-1$^{-/-}$;*dme*-2$^{-/-}$ seedling showing two shoot apex (yellow arrows)

**(D)** *mea*-1$^{-/-}$;*dme*-2$^{-/-}$ seedlings depicting stunted growth, fused cotyledons, phenotype. The scale bar represents **(A-C)** 1mm **(D)** 5 mm.

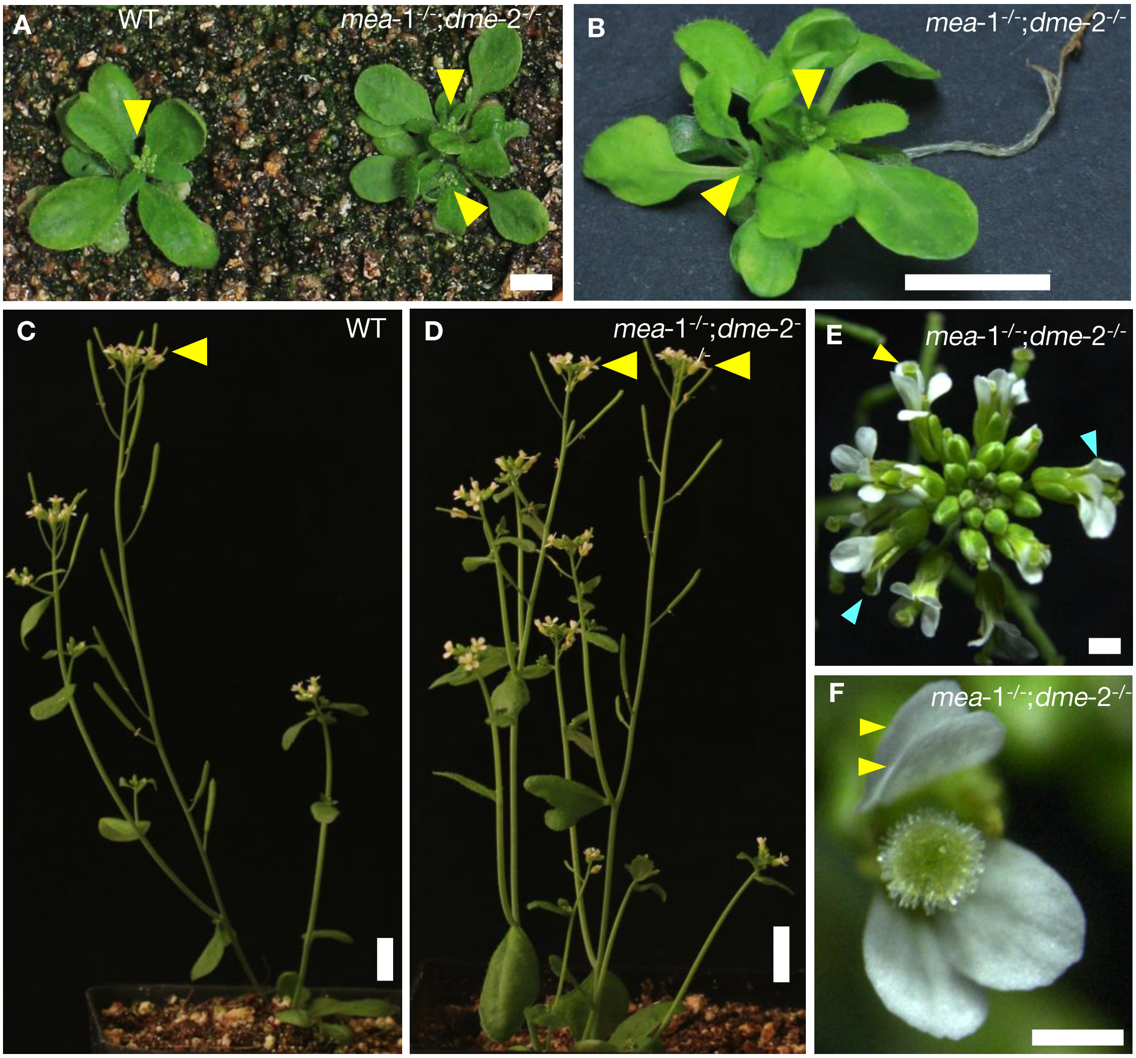


**Supplementary Fig. S3 Phenotypes of *mea*-1$^{-/-}$;*dme*-2$^{-/-}$ flowers and adult plants**

**(A)** *mea*-1$^{-/-}$;*dme*-2$^{-/-}$ adult plant as compared to WT at 10-leaf (15 dpg) stage

**(B)** *mea*-1$^{-/-}$;*dme*-2$^{-/-}$ adult plant depicting ectopic shoot in same plant

**(C)** WT plant showing primary shoot apex (yellow arrow)

**(D)** *mea*-1$^{-/-}$;*dme*-2$^{-/-}$ plant depicting two primary shoot apices (yellow arrow)

**(E)** *mea*-1$^{-/-}$;*dme*-2$^{-/-}$ inflorescence with abnormal floral phenotype such as defective gynoecium (blue arrow), improper petal positioning (yellow arrow)

**(F)** *mea*-1$^{-/-}$;*dme*-2$^{-/-}$ flower showing improper petal positioning. The scale bar represents **(A-D)** 1 cm **(E)** 1 mm and **(F)** 500 μm.

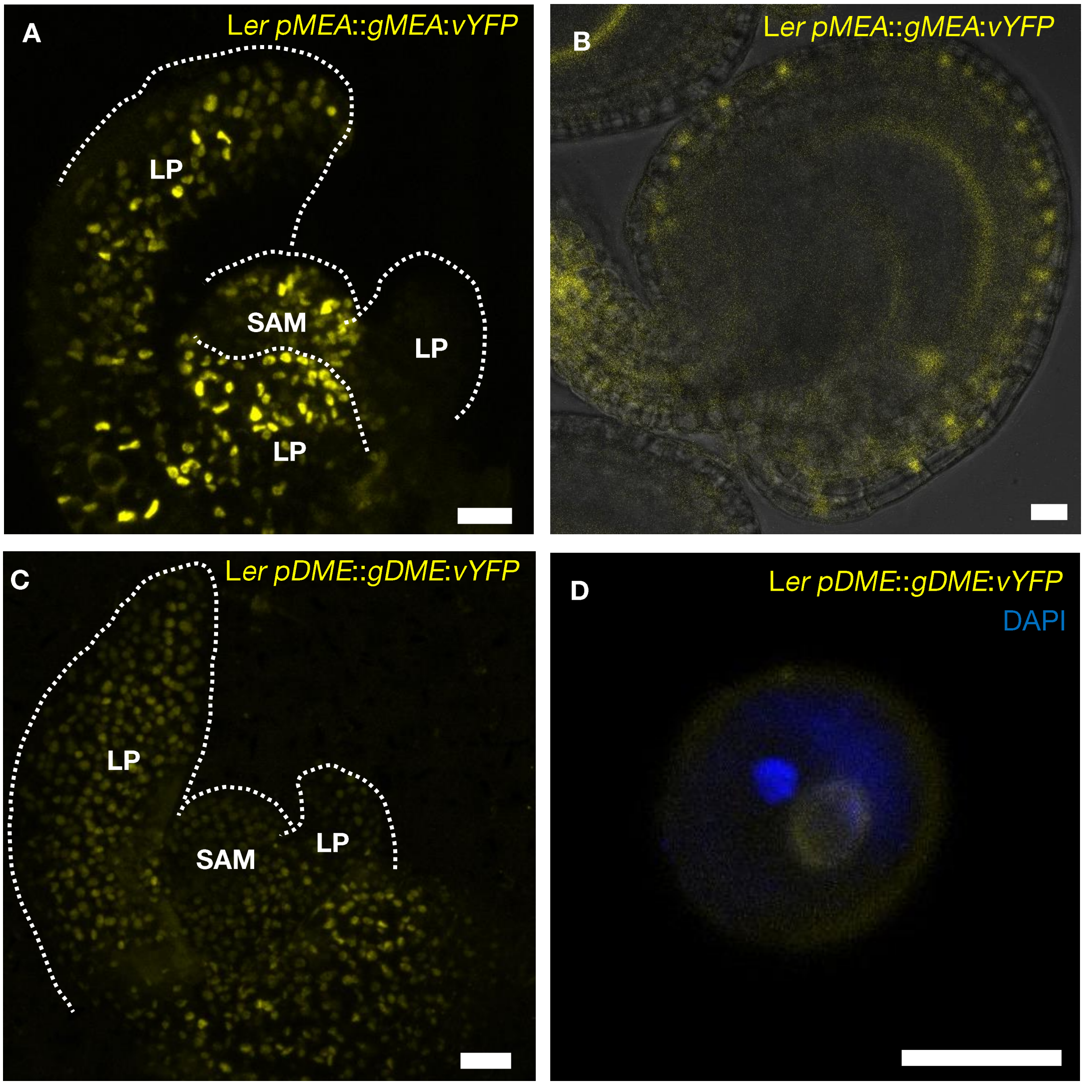


**Supplementary Fig. S4 Expression pattern of MEA-vYFP and DME-vYFP in shoot apex**

Representative images of shoot apices, ovule and bicellular pollen grain showing expression (yellow) of **(A-B)** *pMEA::gMEA:vYFP* and **(C-D)** *pDME::gDME:vYFP*. The meristem dome (SAM) and leaf primordia (LP) are labeled and indicated with a white dashed line and nuclei in bicellular pollen are stained with DAPI (blue). The scale bar represents 20 μm.

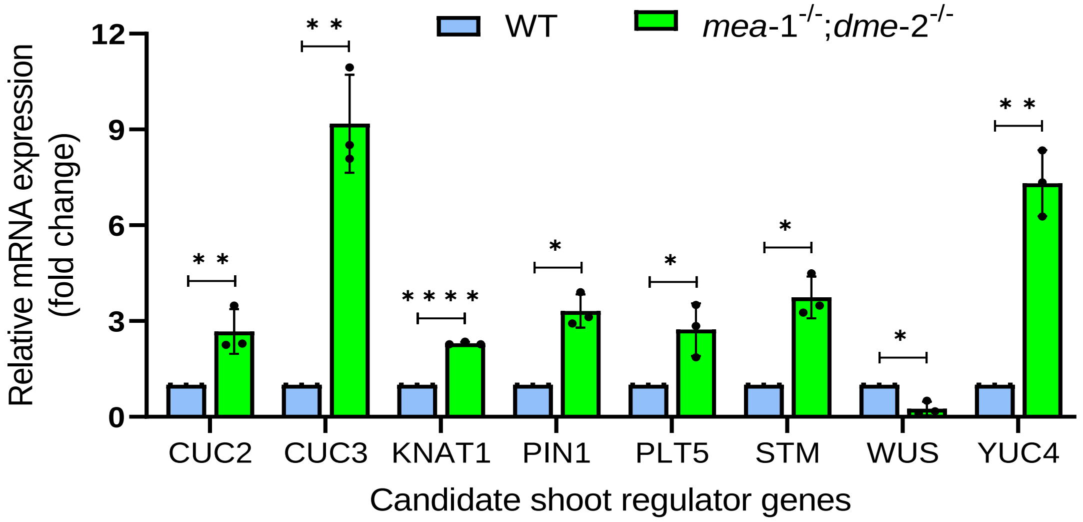


**Supplementary Fig. S5 Expression pattern of shoot regulators in *mea*-1$^{-/-}$;*dme*-2$^{-/-}$ seedlings**

Relative abundance of shoot regulators in 2-3 dpg old seedlings. *CUC2*: 2.52±0.87, **$p$=0.014; *CUC3*: 9.18±1.54, **$p$=0.013; *KNAT1*: 2.30±0.19, ****$p$=0.00002; *PIN1*: 3.31±0.51, *$p$=0.022; *PLT5*: 2.73±0.82, *$p$=0.021; *STM*: 3.74±0.65, *$p$=0.024; *WUS*: 0.26±0.22, *$p$=0.033; *YUC4*: 7.32±1.03, **$p$=0.0081. N=3, statistical test: Multiple unpaired *t*-tests (Holm-Sidak method), with alpha = 0.05. Error bar: SD.

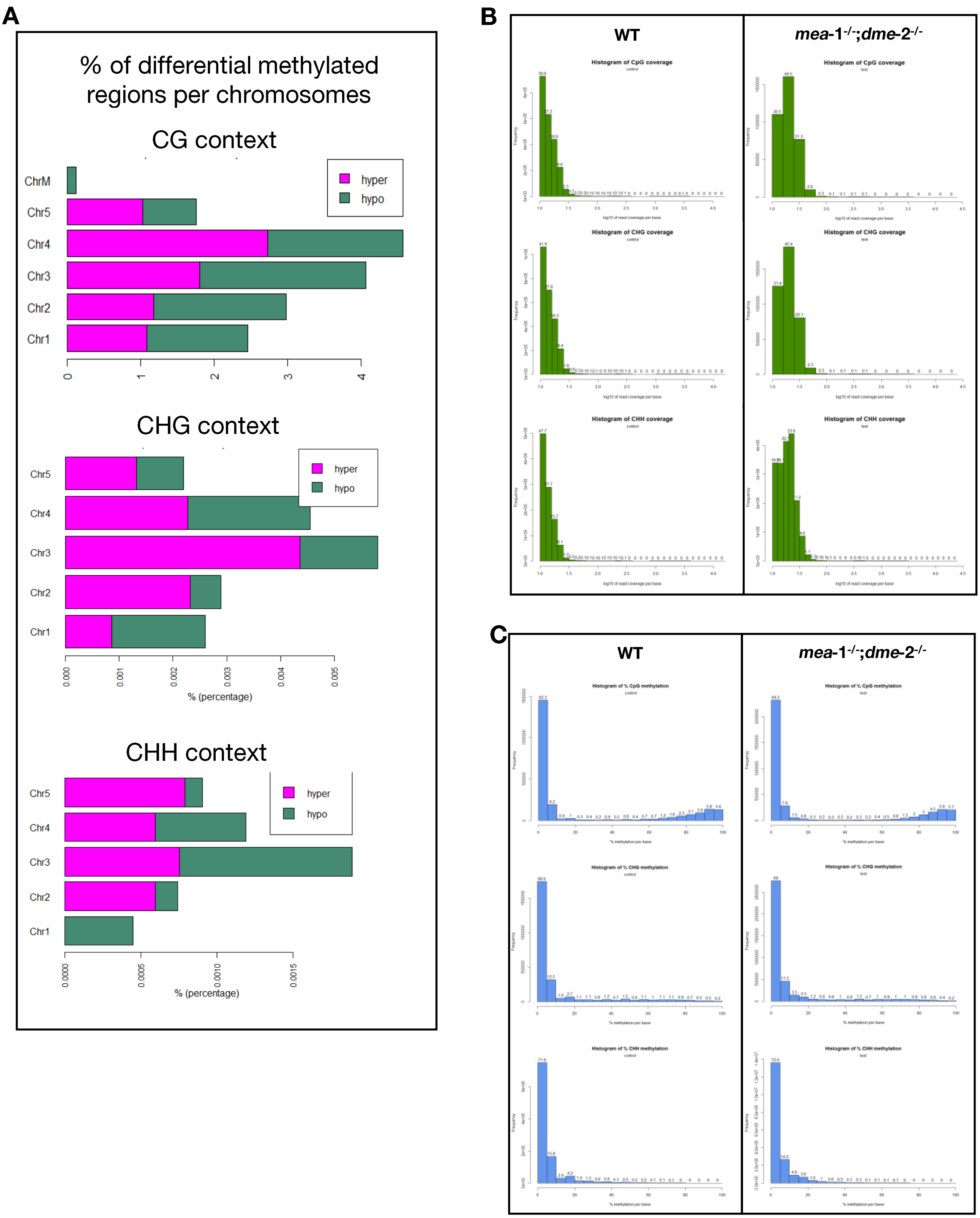


**Supplementary Fig. S6 Bisulfite sequencing analysis**

**(A)** Differential methylation analysis for CG, CHG and CHH context across chromosomes

**(B)** Histogram of CG,CHG and CHH methylation coverage

**(C)** Histogram of % CG, CHG and CHH coverage per base.

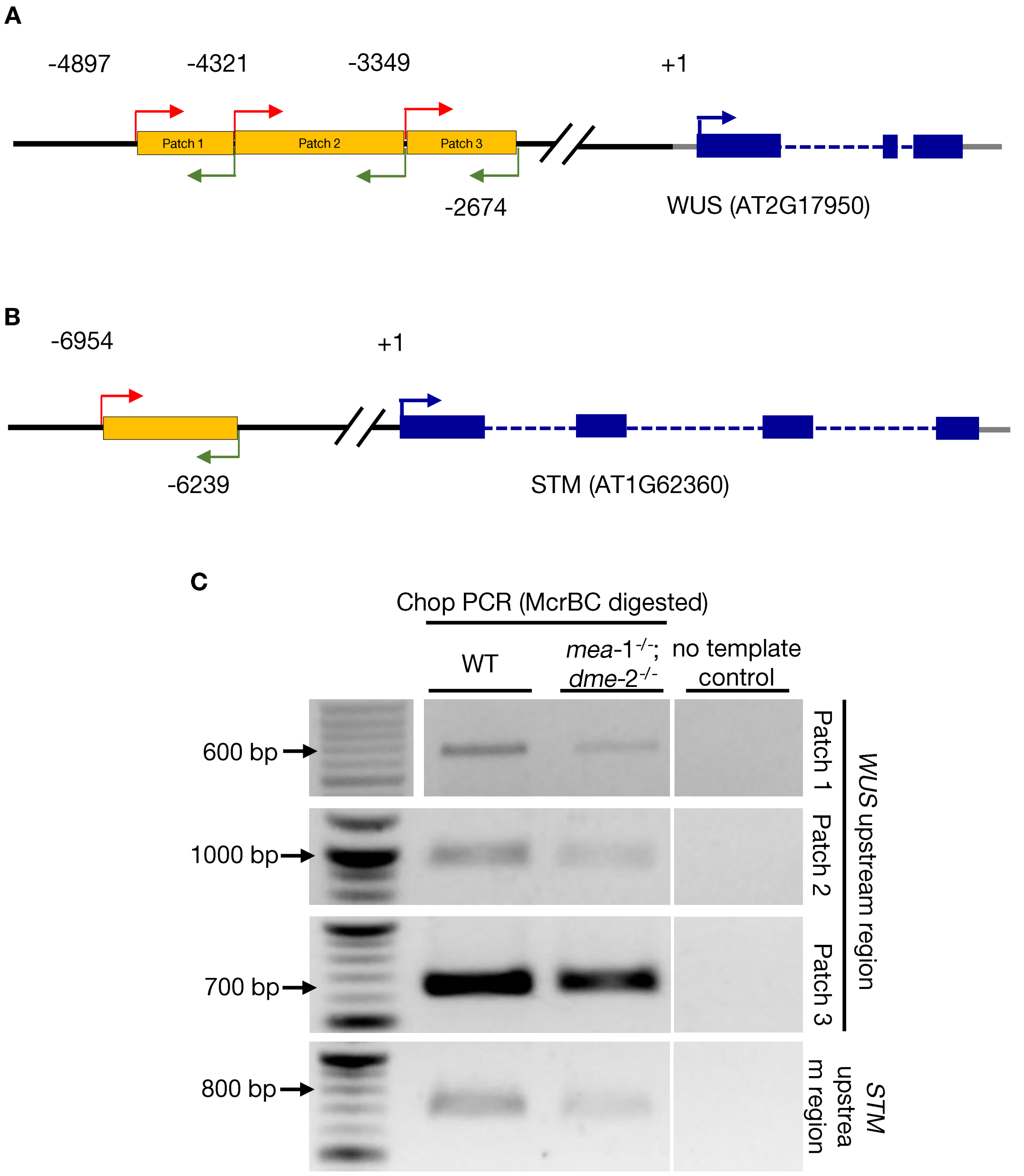


**Supplementary Fig. S7 CHOP PCR for STM and WUS regulatory regions**

**(A-B)** Cartoon representation of *WUS* and *STM* upstream regions showing putative methylated regions (yellow box) and corresponding forward (red) and reverse (green) primers used for Chop PCR

**(C)** Lower amplification in McrBC-chop PCR of hypermethylated regions of *WUS*, and *STM* upstream regions in *mea*-$1^{-/-}$;*dme*-$2^{-/-}$ as compared to hypomethylated WT. Black line: upstream region, gray line: UTR, blue box: exon, blue dotted line: intron.

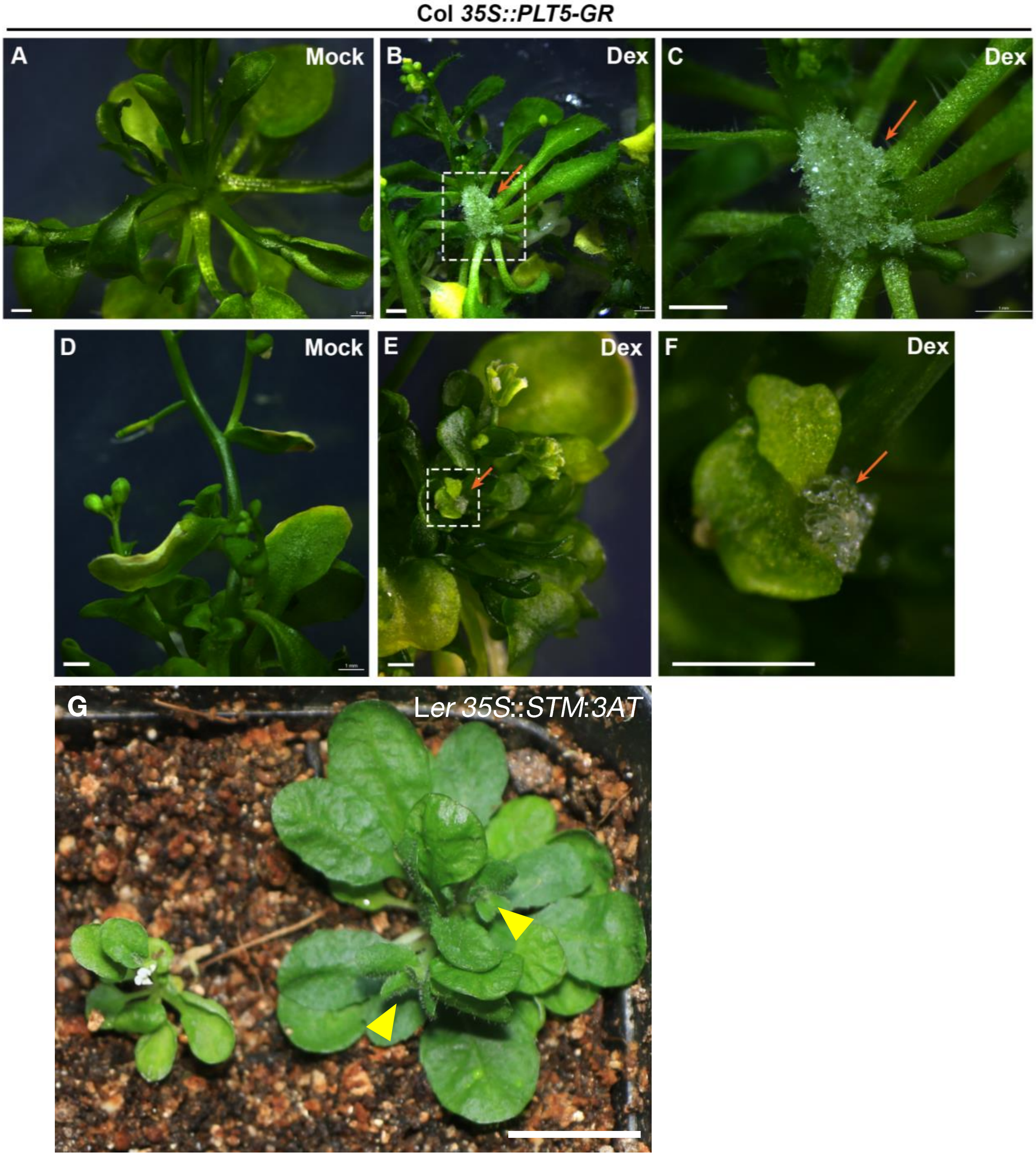


**Supplementary Fig. S8 PLT5 and STM overexpression in seedlings**

**(A-C)** Ectopic cell proliferation from the meristems in the axils of the rosette and

**(D-F)** cauline leaves upon PLT5 overexpression (orange arrows)

**(G)** Ectopic shoots (yellow arrowheads) depicted in STM overexpression

Panels **A,D** represent mock treatment with DMSO. The scale bar represents **(A-F)** 1 mm and **(G)** 1 cm.

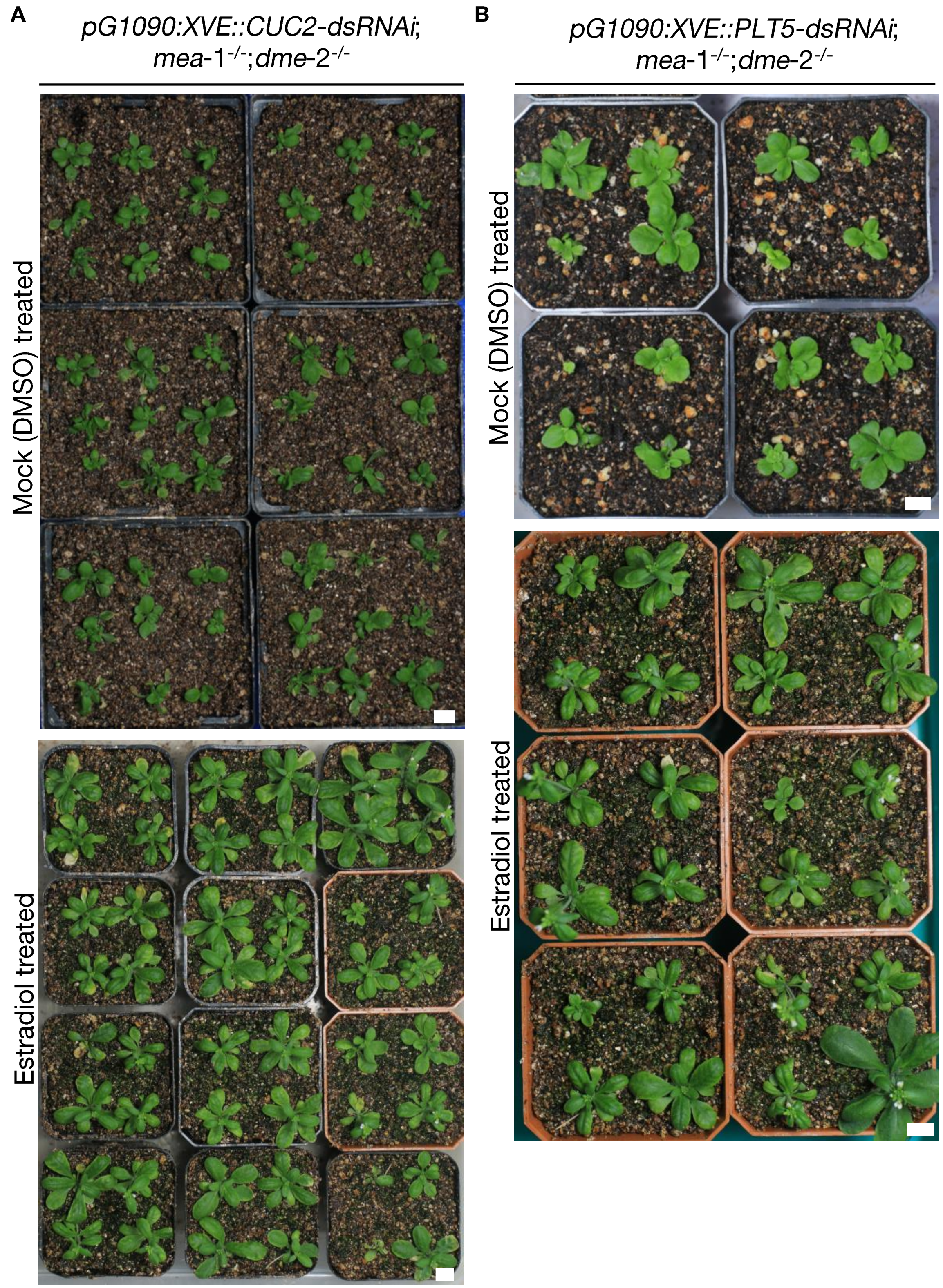


**Supplementary Fig. S9 RNAi mediated downregulation of CUC2 and PLT5 result in rescue of ectopic shoot in *mea*-1$^{-/-}$;*dme*-2$^{-/-}$**

**(A)** *CUC2*-dsRNAi; *mea*-1$^{-/-}$;*dme*-2$^{-/-}$

**(B)** *PLT5*-dsRNAi; *mea*-1$^{-/-}$;*dme*-2$^{-/-}$ plants. The scale bar represents 1 cm.

### SI Material and Methods

**Chop-PCR assay.** Genomic DNA from 2 to 3 dpg old WT and *mea*-1$^{-/-}$;*dme*-2$^{-/-}$ seedlings was isolated using the standard CTAB method. One microgram of DNA was digested with methylation-sensitive restriction enzyme McrBC (NEB, Cat# M0272) at a concentration of 10 units/µg of DNA template in a 30 µl reaction mix, as per manufacturer's instruction, for at least 6 h. After heat inactivation, 4 µl of the digested DNA was used as a template in a 20 µl reaction volume for PCR for 25 cycles.

**Chromosome dosage analysis.** To analyze gross chromosomal changes, the whole genomic sequence from bisulfite sequencing was partitioned into 100 kb bins, and the read counts in each bin were plotted using custom R scripts. The median and 2x median read coverage were also plotted.

**Plasmid construction, molecular cloning, and plant transformation.** Translation fusion lines of DME (*pDME*::*gDME*:*vYFP*) and MEA (*pMEA*::*gMEA*:*vYFP*) were generated using three fragments multisite recombination gateway cloning system (Invitrogen, Cat# 12538120) where the upstream regulatory sequences of *DME* (5000 bp) and *MEA* (5142 bp) were cloned to drive *DME* (8303 bp) and *MEA* (4199 bp) genomic sequences fused with *vYFP*. To constitutively express STM (*35S*::*STM*:*3AT*), 1363 bp CDS region of STM was amplified from *cDNA* using region-specific oligonucleotides and cloned under *35S* constitutive promoter. The entry clones were combined in *pH7m24Gw* based destination vector using multisite recombination gateway cloning system (Invitrogen, Cat# 12538120). Transgenic lines harboring the inducible *35S*::*PLT5*:*GR* construct was describe previously (Prasad et al. 2011). These constructs were introduced into *Agrobacterium tumefaciens* strain C58 by electroporation. Stable transgenic plants were generated by floral-dip method (Clough and Bent 1998).

**Quantitative Real Time-PCR.** For expression analysis of selective shoot regulators and somatic epigenetic factors, we harvested shoot apices from 2 to 3-day-old WT and *mea*-$1^{-/-}$;*dme*-$2^{-/-}$ seedlings. Total RNA was isolated using TRIzol™ reagent as per the manufacturer's instructions. The extracted RNA was treated with NEB(New England Biolabs) DNase I (RNase free) (M0303) according to NEB guidelines to remove the genomic DNA contamination. One microgram of total RNA was used for complementary DNA (cDNA) synthesis by PrimeScript first strand cDNA synthesis kit (TAKARA-Bio, Cat# 6110 ) using an oligo(dT) and random hexamer as oligonucleotides. qPCRs were performed on Mastercycler BIORAD CFX96™ using the oligonucleotides mentioned in Supplementary Table 5. *ACTIN2* (*ACT2*) served as a reference gene control. The reactions were carried out using TB Green® Premix Ex Taq™ II (TAKARA-Bio, Cat no. RR82WR) and incubated at 95 °C for 2 minutes, followed by 40 cycles of 95 °C for 15 s and 60 °C for 30 s. PCR specificity was checked by melting curve analysis. *ACT2* was used for the normalization for all the reactions. At least three independent biological samples were analyzed for each time point mentioned above, and qPCR reactions were set up with three technical replicates for each biological replica. mRNA abundance of target genes was analyzed (Supplementary Table 4) using the normalized $2^{-\Delta\Delta Ct}$ (cycle threshold) method (Livak and Schmittgen, 2001).

**References**


Clough SJ, Bent AF (1998) Floral dip: a simplified method for Agrobacterium-mediated transformation of Arabidopsis thaliana. Plant J 16:735–743. https://doi.org/10.1046/j.1365-313x.1998.00343.x

Livak KJ, Schmittgen TD (2001) Analysis of Relative Gene Expression Data Using Real-Time Quantitative PCR and the 2−ΔΔCT Method. Methods 25:402–408. https://doi.org/10.1006/meth.2001.1262

Prasad K, Grigg SP, Barkoulas M, et al (2011) Arabidopsis PLETHORA transcription factors control phyllotaxis. Curr Biol 21:1123–1128. https://doi.org/10.1016/j.cub.2011.05.009
